# Absence of nascent peptides triggers nonfunctional ribosome decay

**DOI:** 10.64898/2026.05.19.726435

**Authors:** Tomoko Sakata, Kotaro Fujii, Makoto Kitabatake

## Abstract

The nonfunctional rRNA decay (NRD) pathway eliminates defective ribosomes to maintain protein synthesis integrity. Although the Mms1-Rtt101 E3 ligase is known to trigger 25S NRD to clear defective 60S subunits, the specific molecular marks and targets it recognizes remain unknown. Here, we demonstrate that the late-stage maturation factor Reh1 functions as a specific molecular sensor for these defects. While Reh1 is normally displaced from the polypeptide exit tunnel (PET) during successful translation initiation, it is persistently retained on nonfunctional 80S ribosomes incapable of peptide bond formation. This prolonged retention, driven by the absence of a nascent peptide, provides a physical mark that recruits the Mms1 complex via Reh1’s N-terminal domain, triggering site-specific ubiquitination of ribosomal protein Rpl19 (uL6). Our findings reveal that the cell repurposes a maturation factor as a sentinel to monitor translation competence, achieving robust detection of diverse ribosomal and maturation defects through a single molecular pathway.

## Introduction

The ribosome is the macromolecular machine responsible for all cellular protein synthesis, rendering its integrity a fundamental requirement for cellular fitness^1–5^. Despite its structural conservation, the immense scale and complexity of ribosome biogenesis inherently introduce functional heterogeneity, producing defective subunits through biosynthetic errors, assembly failures, or environmental damage^6–9^. Because the catalytic core of the ribosome is mostly RNA-based, even a single point mutation within its key functional domains, such as the decoding center (DC) in the 40S subunit or the peptidyl transferase center (PTC) in the 60S subunit, is sufficient to leave the entire particle translationally inactive. To counter this vulnerability, cells selectively recognize and eliminate these catalytically inert ribosomes through a specialized quality control pathway termed nonfunctional rRNA decay (NRD) ^10–14^.

In the case of the small 40S subunit harboring mutations in the 18S rRNA (termed 18S NRD), recent breakthroughs have elucidated the precise molecular basis of substrate recognition^15–20^. 18S NRD is triggered by the prolonged stalling of a defective 40S subunit harboring an empty A-site as an 80S ribosome at the start codon^12,18^. In mammals, this empty A-site is sensed by the GCN1-GCN2 complex, leading to GCN2 activation and the subsequent induction of the integrated stress response (ISR)^21,22^ to transiently suppress global translation initiation^18^. This translational suppression maintains the stalled ribosome as an isolated monosome, allowing the E3 ligase RNF10^23–25^ (or Mag2 via direct stall-sensing in yeast^17^) to precisely access and ubiquitinate the ribosomal protein uS3, thereby flagging the nonfunctional subunit for downstream clearance^17,18,26^.

In contrast to the 18S NRD pathway, the surveillance mechanism for the large 60S subunit (25S NRD) represents a distinct and largely unresolved challenge. We previously reported that the degradation of nonfunctional 25S rRNA (25S NRD) in *Saccharomyces cerevisiae* requires an E3 ubiquitin ligase complex containing Mms1, Rtt101, and Crt10^13,27^. Following Mms1-mediated ubiquitination, the defective 60S subunit within the 80S particle is recognized by the Cdc48-Ufd1-Npl4 AAA-ATPase complex, dissociated from the 40S subunit, and subsequently degraded by the proteasome^28^. While this downstream degradation machinery has been well-characterized, the upstream mechanism of substrate recognition—how the Mms1 complex initially distinguishes the dysfunctional state from the functional one—has remained elusive.

In this study, we identify the late-stage ribosome maturation factor Reh1^29,30^ as a physical interaction partner of the Mms1 E3 ligase complex. We found that Reh1 is selectively enriched on 80S ribosomes carrying nonfunctional 25S rRNA compared to those with wild-type 25S rRNA. This discovery reveals a mechanism wherein the absence of a nascent peptide causes the prolonged retention of Reh1 within the polypeptide exit tunnel (PET), thereby marking defective ribosomes for E3 recruitment. Furthermore, we demonstrate that this retention precisely positions the E3 complex to ubiquitinate the 60S protein Rpl19 (uL6) at its 40S-contacting protrusion, providing a strategic structural hallmark for the subsequent Cdc48-mediated disassembly of nonfunctional 80S particles. Our findings reveal that the cell repurposes a maturation factor as a sentinel to monitor translation competence, linking PET vacancy directly to the initiation of 25S NRD.

## Results

### Diverse nonfunctional mutant 25S rRNAs trigger Mms1-dependent degradation

To explore the sensing mechanism of 25S NRD, we first addressed the limited repertoire of available substrates. Previous studies have relied on a few specific mutations within the peptidyl transferase center (PTC), such as A2451U or U2585A^12–14,27,28^. Thus, it remains unclear whether the Mms1 complex merely recognizes a localized structural perturbation unique to these PTC mutations, or if it senses a more universal molecular signature of 60S dysfunction. To distinguish between these possibilities, we expanded the repertoire of defective ribosomal variants.

We assessed the functionality of various 25S rRNA mutants using a yeast strain harboring a temperature-sensitive endogenous RNA polymerase I^31^. In this system, cell growth at the restrictive temperature is strictly dependent on the functionality of the plasmid-encoded 25S rRNA transcribed by RNA polymerase II^32^ (Figure 1A). In addition to the previously characterized mutations in PTC, we identified a broad spectrum of 25S rRNA mutations that result in functional insufficiency. While representative examples are shown in Figure 1B, the comprehensive results for all identified variants are detailed in Figure S1A. While many of these mutations are centered around the PTC, they also distribute to more distant regions of the 25S rRNA (Figure1B). Remarkably, northern blot time-course analysis following transcriptional shut-off of plasmid-encoded rRNAs revealed that the majority of these nonfunctional variants were unstable in wild-type cells but were significantly stabilized upon the deletion of *MMS1* (Figure 1B and C). Together, these results demonstrate that the 25S NRD pathway is capable of targeting a much broader range of ribosomal defects than previously appreciated, irrespective of their precise location or the nature of the nucleotide substitution (Figure 1C and Figure S1C). This broad target spectrum firmly establishes that the Mms1 complex does not merely recognize a specific local structural perturbation, but rather responds to a more universal signature of ribosomal dysfunction.

**Figure 1.**
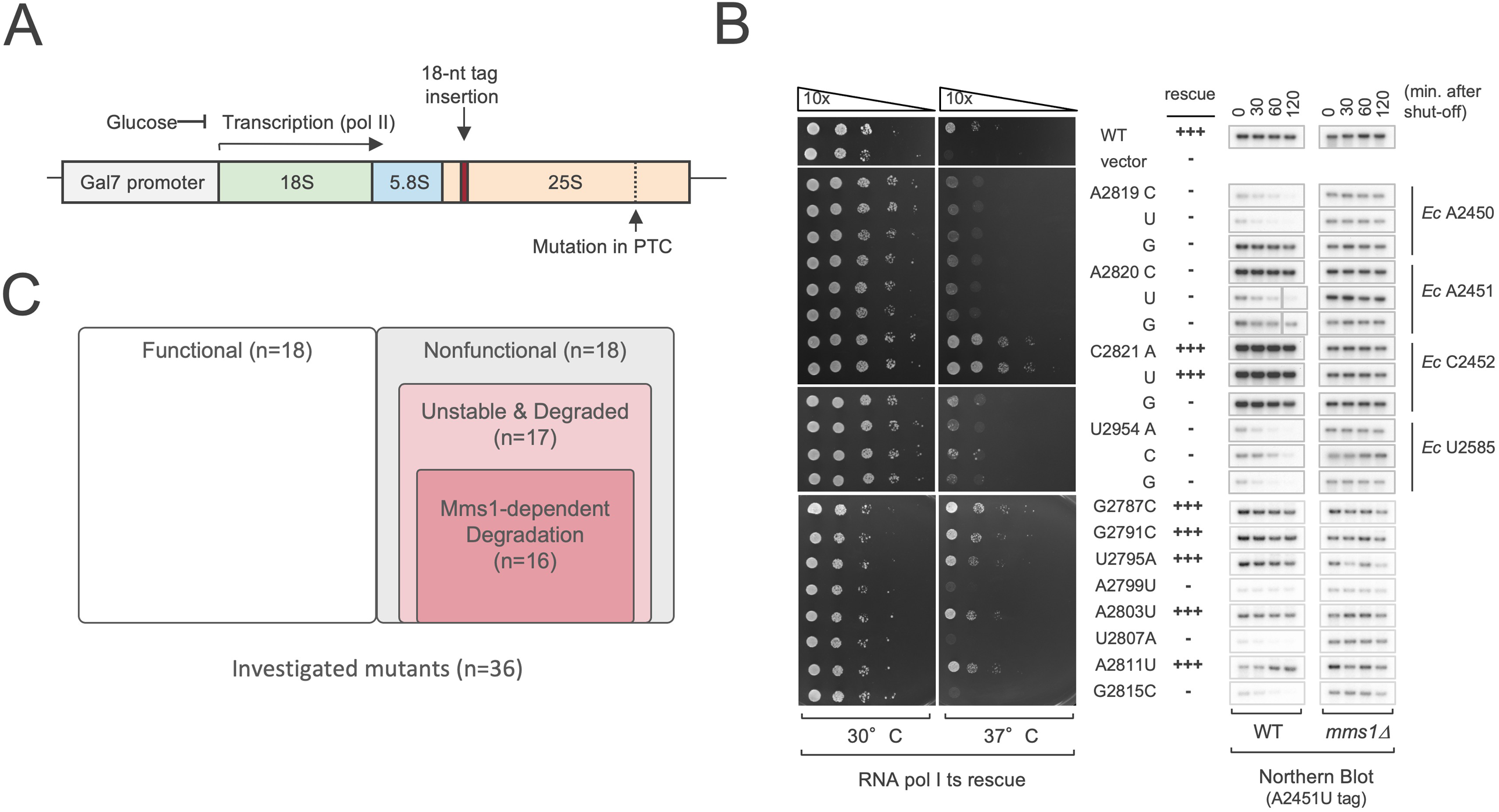
Mms1-dependent 25S NRD universally targets diverse nonfunctional 60S subunits. **(A) Schematic of the 25S rRNA reporter system.** The yeast 35S pre-rRNA is transcribed from the *GAL7* promoter by RNA polymerase II. A short 18-nt tag inserted into a non-conserved loop of the 25S rRNA serves as a marker for Northern blot and qRT-PCR detection. **(B) Functional and stability analysis of 25S rRNA mutants.** Left: Colony formation assay of the indicated 25S rRNA mutants (middle) expressed in an RNA polymerase I temperature-sensitive strain. Right: Northern blot analysis of the tagged 25S rRNA variants following transcriptional shut-off in wild-type (WT) or *mms1*Δ strains. **(C) Venn diagram of 25S rRNA variant classifications.** Of the 36 tested variants, 18 are functional and 18 are nonfunctional. Within the nonfunctional group, 17 are categorized as unstable and degraded, 16 of which undergo Mms1-dependent degradation, exhibiting at least partial stabilization in the *mms1*Δ strain. The complete dataset, including individual variant assignments, is presented in Figure S1.

### Reh1 acts as a specific sensor bridging the E3 ligase complex and non-functional 80S ribosomes

We initially hypothesized that this detection might rely on known surveillance pathways, such as the Ribosome-associated Quality Control (RQC) system^11^. However, 25S NRD remained fully functional in strains lacking core RQC factors^33,34^ (*hel2*Δ, *rqc1*Δ, and *rqc2*Δ), as well as in those lacking major ribosome-associated chaperones^35–38^ or ubiquitin regulators^39–41^ (Figure S2A, B). These results indicate that 25S NRD operates independently of established quality control pathways, prompting us to search for a distinct, highly specialized sensor.

Reasoning that such a sensor must physically bridge the defective ribosome to the degradation machinery, we analyzed physical interactions between the Mms1-Rtt101-Crt10 E3 complex and several candidate ribosome-associated factors^42–44^. Our biochemical pull-down analysis revealed that Reh1 exhibits a strong and direct affinity for the E3 complex (Figure 2A). Reh1 is a structural homolog of Rei1^29,37,45,46^, and both factors are known to insert their C-terminal tails into the PET during late-stage 60S maturation^30,45,47^. While Rei1’s prior insertion is essential for pre-60S biogenesis^45^, the functional significance of Reh1’s subsequent tunnel occupancy had remained unclear^30^. Mutational mapping further identified the N-terminal zinc-finger domain of Reh1 as the essential platform for this physical interaction (Figure 2B).

**Figure 2.**
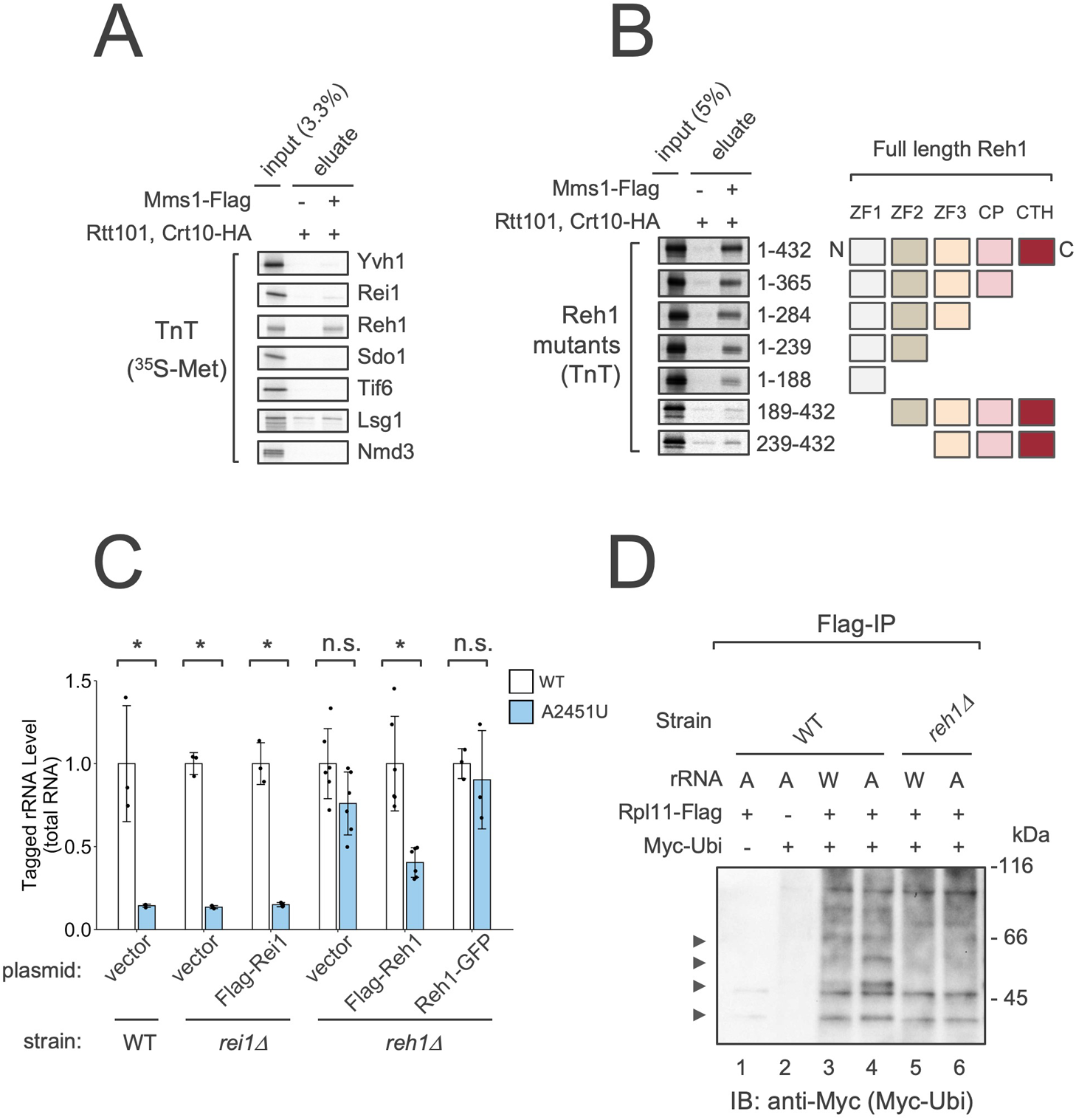
Reh1 is the molecular sensor for 25S NRD required for ribosome ubiquitination. **(A) In vitro binding of candidate factors to the Mms1 E3 complex.** Left: 35S-labeled in vitro-translated inputs. Middle: In vitro binding to Flag-IP fractions from a strain expressing Rtt101/Crt10-HA without Mms1-Flag (nonspecific control). Right: In vitro binding to the purified Mms1-Flag/Rtt101/Crt10-HA complex. **(B) Mapping of the Reh1 domain interacting with the Mms1 E3 complex.** In vitro binding assays using 35S-labeled Reh1 truncation mutants. Amino acid coordinates of each fragment (from the 432-aa full-length protein) are indicated to the right of the gel, aligned with a schematic of the corresponding regions within the full-length domain architecture on the far right. **(C) Requirement of Reh1 for 25S NRD.** The 25S NRD reporter (wild-type or A2451U) was expressed in the indicated strains (WT, *rei1*Δ, or *reh1*Δ) transformed with various plasmids. Tagged reporter RNA levels in log-phase cells were quantified by qRT-PCR to assess whether Reh1 is required for 25S NRD, compared with its paralog Rei1. **(D) Ribosome ubiquitination in WT and *reh1***Δ **strains.** Ribosomes were isolated by Flag-IP from WT or *reh1*Δ strains co-expressing Rpl11-Flag, Myc-ubiquitin, and wild-type (W) or nonfunctional A2451U (A) 25S rRNA from three plasmids. Ubiquitinated ribosomal products were visualized by anti-Myc immunoblotting.

The *in vivo* functional significance of Reh1 was confirmed by the observation that its deletion completely stabilized 25S NRD substrates (Figure 2C). In contrast, deletion of its homolog, *REI1*, had no effect on substrate stability. Introduction of a Flag-tagged Reh1 plasmid partially rescued the degradation defect in *reh1*Δ cells; the incomplete recovery was likely due to minor steric interference from the N-terminal Flag tag (Figure 2C). Crucially, this substrate stabilization correlated with a total loss of selective ubiquitination of nonfunctional ribosomes by the Mms1 E3 ligase in *reh1*Δ cells (Figure 2D, lane 6), establishing Reh1 as an indispensable factor for Mms1-mediated target recognition.

To determine the molecular basis of this specificity, we generated domain-swap chimeras between Reh1 and Rei1 (Figure S3A). In *reh1*Δ cells, the Reh1N-Rei1C chimera successfully restored 25S NRD, whereas the reciprocal Rei1N-Reh1C chimera failed to do so (Figure S3B). Conversely, the Rei1N-Reh1C variant successfully rescued the cold-sensitive phenotype of *rei1*Δ cells (Figure S3C). These results demonstrate that the N-terminal E3-binding domain—which we identified as the physical interface with Mms1 in the previous section—is the critical determinant specifying Reh1 function in 25S NRD, whereas the C-terminal tail is functionally interchangeable between the two homologs for this process.

The importance of the C-terminus was further supported by the observation that a Reh1 variant carrying a bulky C-terminal GFP tag (Reh1-GFP) failed to rescue the NRD defect (Figure 2C). Consistently, a C-terminally truncated variant (Flag-Reh1ΔC) similarly failed to restore NRD activity (Figure S3B), highlighting that an unobstructed C-terminus is vital for its sensing role. Unlike many pre-60S biogenesis factors, the absence of Reh1 did not result in significant growth defects or gross alterations in polysome profiles (Figure S3D), indicating that its primary role is specialized for quality control rather than general 60S maturation.

To determine the exact translational stage at which Reh1 monitors ribosome functionality, we next analyzed the distribution of the stabilized nonfunctional rRNAs. Interestingly, the nonfunctional 25S rRNAs in *reh1*Δ cells predominantly accumulated within the 80S fraction (Figure 3A), in contrast to the distribution of nonfunctional 25S rRNAs in 60S fraction in a wild-type strain. Considering that Reh1 joins the pre-60S subunit at a late maturation stage^47^ and is typically released during the transition to active translation^30^, this 80S accumulation strongly suggests that 25S NRD primarily targets ribosomes in the 80S state. This finding led us to hypothesize that Reh1 acts as the specific molecular sensor by selectively recognizing and remaining associated with nonfunctional 80S ribosomes. To test this *in vivo* binding selectivity, we performed immunoprecipitation of Reh1 from cell lysates, which revealed a striking preferential enrichment of nonfunctional mutant 25S rRNAs over their wild-type counterparts (Figure 3B).

**Figure 3.**
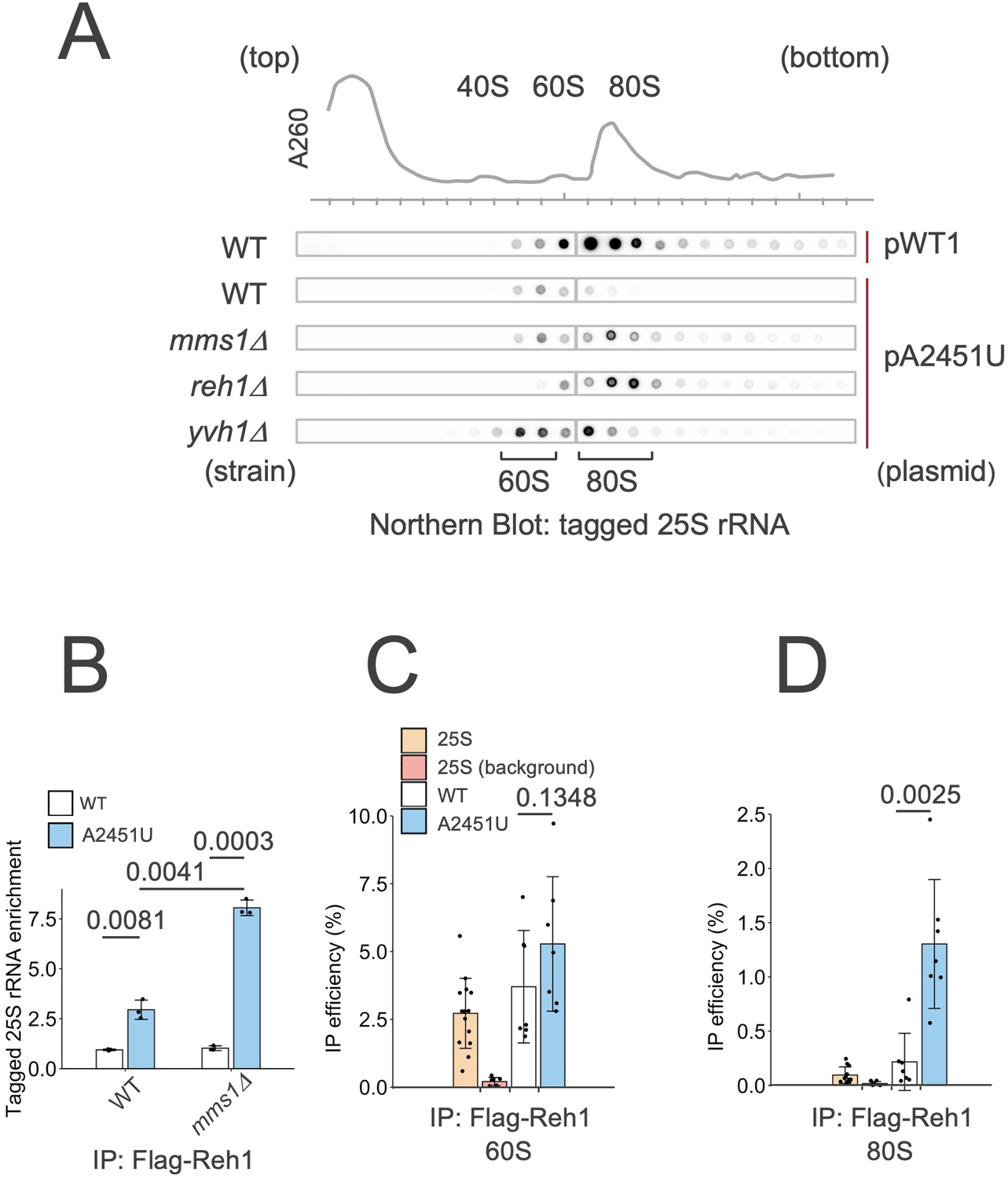
Selective retention of Reh1 on nonfunctional 80S ribosomes serves as a molecular beacon. **(A) Sucrose gradient sedimentation of 25S NRD reporters.** A representative A260 absorbance profile is aligned at the top to correlate gradient fractions with ribosome peaks. The wild-type (pWT1) or nonfunctional (pA2451U) 25S NRD reporter was expressed in the indicated strains (WT, *mms1*Δ, *reh1*Δ, or *yvh1*Δ), and the distribution of the tagged reporter RNAs across the fractions was analyzed by Northern blotting. **(B) Preferential enrichment of nonfunctional 25S rRNAs on Reh1.** qRT-PCR analysis of copurified wild-type (WT) and nonfunctional (A2451U) 25S rRNAs following Flag-Reh1 immunoprecipitation (IP) from WT or *mms1*Δ total cell lysates. **(C and D) Ribosome-state-specific selectivity of Reh1 binding.** Flag-Reh1 was immunoprecipitated from sucrose density gradient-separated 60S (C) or 80S (D) fractions, followed by qRT-PCR analysis of copurified rRNAs. Bar labels indicate the targets detected: total 25S rRNA (endogenous and reporter; “25S”), tagged wild-type (“WT”), or nonfunctional (“A2451U”) reporters. “25S (background)” denotes total 25S rRNA recovered from a mock IP using a strain lacking Flag-Reh1 expression.

Crucially, we discovered that this discriminatory binding is strictly specific to the 80S state. When analyzing 60S and 80S fractions separately, we found that Reh1 associated with both functional and nonfunctional subunits in the 60S fraction without clear distinction (Figure 3C). In sharp contrast, within the 80S fraction, Reh1 was exclusively retained on nonfunctional particles, whereas it was absent from functional 80S ribosomes (Figure 3D). This selective retention within the 80S complex represents the first identified physical hallmark of a nonfunctional ribosome. By persistently occupying a stage from which it should normally be evicted, Reh1 serves as a molecular beacon that flags defective particles for the Mms1 machinery.

### The Mms1 complex specifically targets the C-terminus of Rpl19 for ubiquitination

Having identified Reh1 as the molecular beacon that flags defective ribosomes, we next sought to determine the direct protein substrate of the Mms1-Rtt101 complex. We performed a proteomic screen using cells co-expressing His-Myc-tagged ubiquitin^48,49^ and nonfunctional 25S rRNA^13^. Following the enrichment of ubiquitinated proteins under denaturing conditions (Figures S4A, B). mass spectrometry identified 11 candidate proteins (Figure S4C). To rigorously isolate covalently modified targets, we systematically validated these candidates through individual biochemical assays by monitoring the poyubiquitin ladder of each HA-tagged protein (Figure S4D). This approach identified Rpl19 (uL6) as the sole target whose ubiquitination was strictly dependent on Mms1 (Figure 4A).

**Figure 4.**
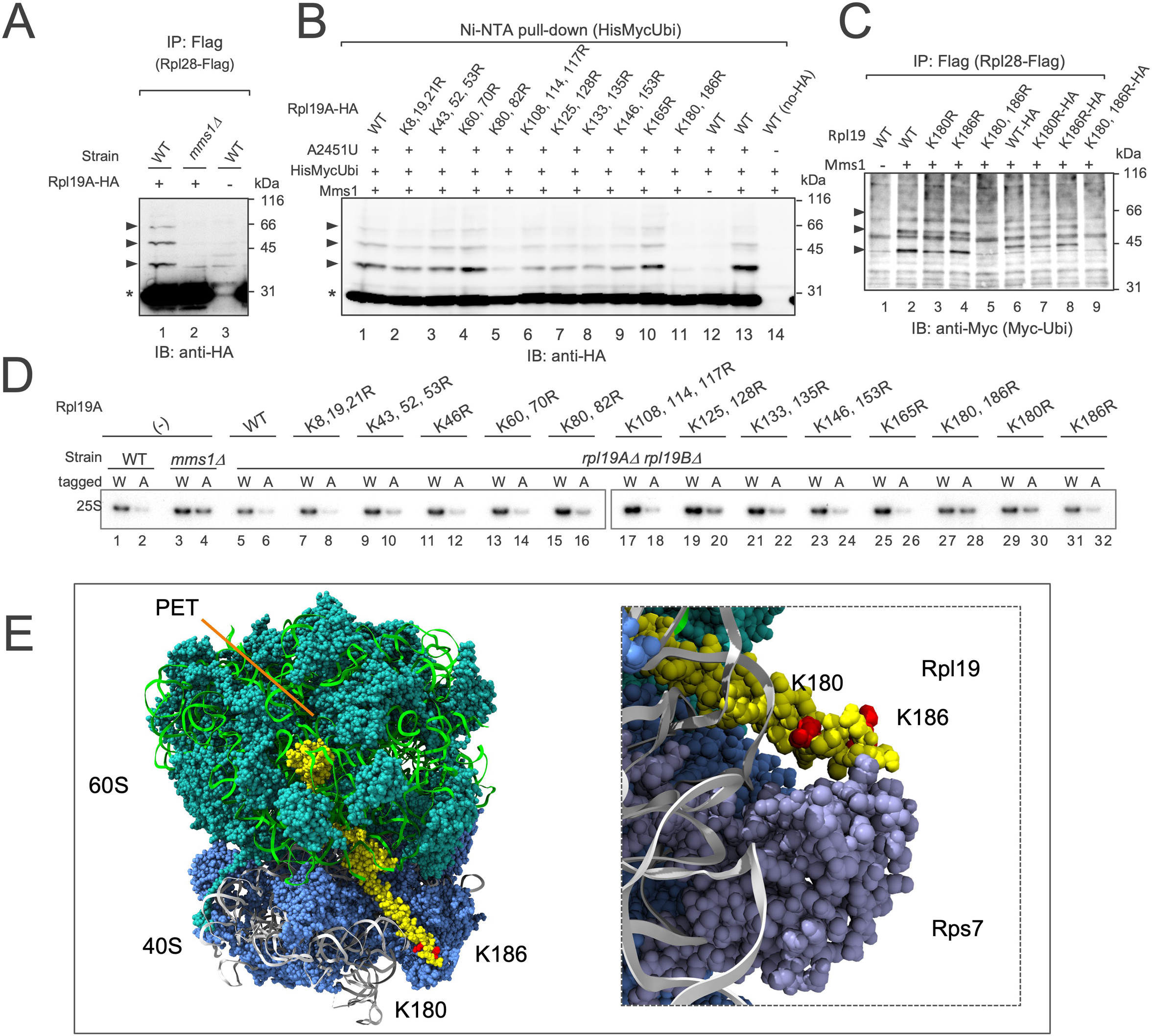
Site-specific ubiquitination of Rpl19 (uL6) by Mms1 triggers 25S NRD. **(A) Mms1-dependent ubiquitination of Rpl19.** Ribosomes were isolated via Rpl28-Flag IP from WT or *mms1*Δ strains expressing Rpl19A-HA. Anti-HA immunoblotting visualizes unmodified Rpl19A-HA (asterisk) and a polyubiquitinated ladder (black arrowheads). This ubiquitinated signal is Mms1-dependent (lane 1 vs. lane 2) and specific to Rpl19A-HA expression (lane 3). **(B) Identification of the Rpl19 ubiquitination site.** Denaturing pull-down of His-Myc-ubiquitin followed by anti-HA immunoblotting from cells expressing the indicated Rpl19A-HA lysine-to-arginine (K-to-R) mutants. Symbols are as in (A). **(C) Total ribosomal ubiquitination in Rpl19 K-to-R mutants.** Ribosomes purified via Rpl28-Flag IP were analyzed by anti-Myc immunoblotting. Lanes 2–5 and lanes 6–9 express untagged and C-terminally HA-tagged versions of the indicated Rpl19 mutants, respectively, with the observed mobility shift reflecting the presence of the HA tag. **(D) Effect of Rpl19 mutations on 25S NRD.** Northern blot analysis of steady-state 25S NRD reporter RNAs (wild-type [W] or nonfunctional A2451U [A]). Reporters were expressed in *rpl19a*Δ *rpl19b*Δ strains sustained exclusively by a plasmid encoding wild-type or the indicated Rpl19 mutants (see Figure S5 for strain construction via plasmid shuffling). **(E) Spatial arrangement of Rpl19 ubiquitination sites.** Three-dimensional model derived from PDB: 4V88, showing the locations of the identified ubiquitination sites (K180 and K186, red) on Rpl19 (yellow) relative to the 60S subunit (green), the 40S subunit (blue), and the polypeptide exit tunnel (PET). The inset provides a magnified view of the ubiquitination sites, demonstrating that this region serves as the contact interface with Rps7 (lavender) of the 40S subunit.

To pinpoint the exact modification sites, we generated a comprehensive set of Rpl19 variants covering all lysine residues, mutated either individually or in small clusters. These variants were introduced into yeast using a plasmid shuffling strategy to replace the endogenous genes (Figures S5A, B). While single or clustered mutations across most of the protein failed to abolish the modification, we found that the K180R/K186R double mutation within the C-terminal extension completely eliminated the ubiquitination signal (Figures 4B and C). Individual substitution of either residue did not block ubiquitination (Figure 4C), indicating that the E3 ligase can flexibly modify either of these adjacent residues to ensure the robust ubiquitination of the defective subunit. Importantly, this site-specific ubiquitination is functionally required for 25S NRD, as the Rpl19-K180R/K186R double mutation significantly stabilized nonfunctional 25S rRNA variants (Figure 4D, lane 28).

Intriguingly, these target lysines are situated near the tip of the characteristic C-terminal extension of Rpl19, which protrudes towards the 40S subunit surface (Figure 4E). Given that these target lysines on Rpl19 are located at a distance from the PET where Reh1’s C-terminus is inserted, this finding raises the question of how the rest of the Reh1 molecule spans this structural gap on the ribosomal surface to guide the E3 complex.

### CRAC mapping of the Reh1-mediated E3 recruitment

To bridge the structural gap, we next sought to determine the precise location of the Reh1 N-terminal domain, which directly interacts with the Mms1-Rtt101 complex. Based on previous structural studies, the C-terminal tail of Reh1 is anchored deep within the PET, but its N-terminal “recruiter” domain has remained structurally unresolved^47^. In available cryo-EM maps, this domain is consistently disordered, presumably reflecting the high conformational flexibility required to scan the ribosomal surface and facilitate E3 docking.

To define the *in vivo* ribosomal neighborhood of the Reh1 N-terminus, we performed high-resolution mapping using UV cross-linking and analysis of cDNAs (CRAC)^50^. This analysis identified two prominent cross-linking peaks on the 25S rRNA within Domain III and Domain VI^51^ (Figures S6A, B). Mapping these hits onto the ribosome structure^4^ revealed that both sites cluster on the solvent-exposed 60S surface in close proximity to the ribosomal protein Rpl19 (uL6) (Figure S6C). This structural proximity between the Reh1 N-terminal docking sites and the Rpl19 provides a compelling spatial explanation for substrate targeting. The presence of multiple rRNA contacts suggests that the flexible N-terminal domain of Reh1 docks dynamically around this region, precisely positioning the Mms1 complex near its substrate.

### Reh1 as the universal beacon for the nonfunctional ribosomes in the quality control of pre-60S subunits

Because Reh1 functions during the late, cytoplasmic stages of pre-60S maturation, we investigated whether other factors involved in this late maturation also participate in 25S NRD. Indeed, the nonfunctional 25S rRNA was stabilized in *yvh1*Δ cells harboring an empty vector, which was rescued by wild-type Yvh1 (Figure 5A). Based on this stabilization, we initially expected to see a corresponding loss of target ubiquitination. Strikingly, however, we observed the exact opposite; *yvh1*Δ cells exhibited a massive, constitutive increase in ribosomal protein ubiquitination, even in the absence of nonfunctional 25S rRNA (Figure 5B, lane 1). Importantly, this hyper-ubiquitination remained completely dependent on Reh1 (Figure 5B, lane 2). The apparent stabilization of nonfunctional 25S rRNA in *yvh1*Δ cells is likely because this pervasive, non-selective ubiquitination overwhelms the degradation machinery, effectively masking the system’s ability to specifically identify defective particles.

**Figure 5.**
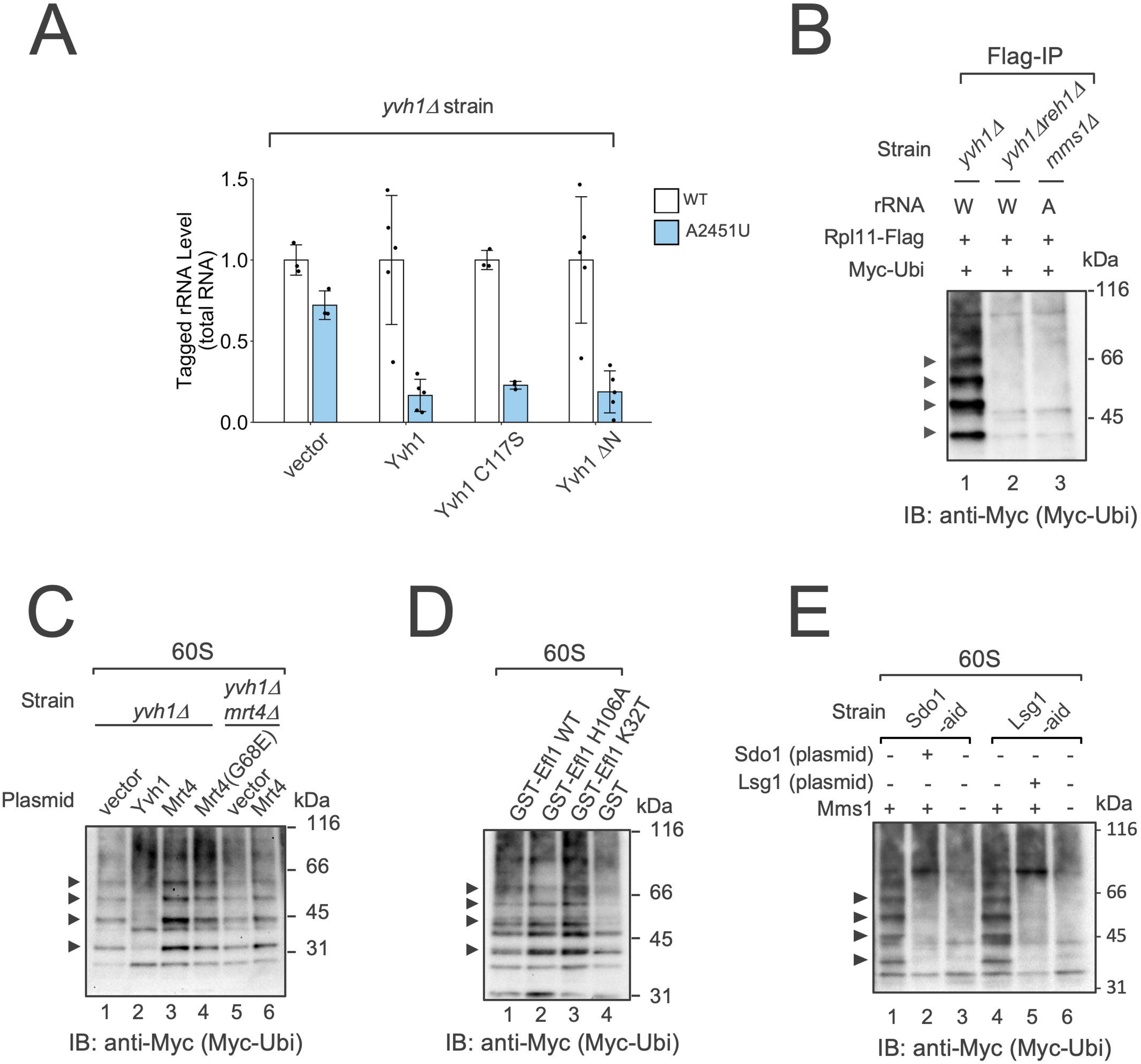
Defects in 60S maturation trigger Reh1-mediated ubiquitination as a universal surveillance response. **(A) Impairment of 25S NRD in *yvh1*Δcells.** qRT-PCR analysis of tagged wild-type or nonfunctional 25S NRD reporter levels in *yvh1*Δ cells. Suppression of 25S NRD activity in the absence of Yvh1 is fully rescued by the expression of wild-type Yvh1 or any of its phosphatase-deficient mutants (C117S and ΔN). **(B) Reh1-dependent ribosomal hyper-ubiquitination induced by Yvh1 loss.** Immunoblot analysis of total ribosomal ubiquitination in *yvh1*Δ, *yvh1*Δ *reh1*Δ, and *mms1*Δ strains. A massive hyper-ubiquitination ladder is exclusively observed in the *yvh1*Δ single mutant, demonstrating its strict dependence on Reh1. **(C) Impact of Mrt4 displacement on 60S ubiquitination.** 60S fractions were isolated via sucrose density gradient centrifugation from *yvh1*Δ or *yvh1*Δ *mrt4*Δ strains expressing the indicated combinations of Yvh1, Mrt4, and Mrt4-G68E. Equal amounts of the recovered 60S subunits were analyzed by anti-Myc immunoblotting to monitor ribosomal ubiquitination levels. **(D) Impact of dominant-negative Efl1 mutations on 60S ubiquitination.** 60S fractions were isolated via sucrose density gradient centrifugation from cells overexpressing the indicated Efl1 variants. Ubiquitination levels of the recovered 60S subunits were analyzed by anti-Myc immunoblotting. **(E) Impact of Sdo1 or Lsg1 depletion on 60S ubiquitination.** 60S fractions were isolated via sucrose density gradient centrifugation from auxin-treated *sdo1-AID* or *lsg1-AID* cells, with or without additional *mms1*Δ mutation. Lanes 2 and 5 express plasmid-encoded Sdo1 or Lsg1 as rescue controls, respectively. Ubiquitination levels of the recovered 60S subunits were analyzed by anti-Myc immunoblotting.

To understand what triggers this hyper-ubiquitination, we performed a domain analysis of Yvh1. During late 60S maturation, Yvh1 facilitates the displacement of Mrt4 to allow P0 stalk assembly^52,53^. It has been reported that the phosphatase domain of Yvh1 is dispensable for pre-60S maturation^52,54,55^. Consistently, we found that neither the enzymatic activity nor the phosphatase domain itself was required to suppress the *yvh1*Δ phenotype, as expression of either a catalytically inactive mutant (C117S) or the N-terminal deletion mutant lacking the entire domain effectively rescued the defect (Figure 5A). This strongly suggests that the Reh1-Mms1 system does not monitor Yvh1’s enzymatic activity, but rather the physical progression of maturation—specifically, the successful displacement of Mrt4.

Supporting this model, overexpression of wild-type Mrt4 further enhanced the ubiquitination signal, whereas expression of the Mrt4-G68E variant^56^, which dissociates spontaneously from the pre-60S subunit even in the absence of Yvh1^52,53^, significantly reduced the ubiquitination level (Figure 5C). Furthermore, complete deletion of *MRT4* in the *yvh1*Δ background (*yvh1*Δ *mrt4*Δ) led to an even more pronounced reduction in ubiquitination, suppressing the signal below the level of the G68E variant (Figure 5C). These results demonstrate that the 25S NRD machinery serves as a broader surveillance system that monitors the timely progression of ribosome maturation, triggering an E3-mediated response whenever a maturation stall occurs.

To determine whether this surveillance represents a broader response to defective maturation, we examined other late-stage maturation factors responsible for the final translational activation of the 60S subunit. Consistent with our observations in *yvh1*Δ cells, hyper-ubiquitination was also triggered by the expression of a dominant-negative Efl1 GTPase mutant, which blocks the evacuation of early maturation factors^57^ (Figure 5D). We confirmed that this condition indeed induced the expected pre-60S maturation stall (Figure S7A). Similarly, hyper-ubiquitination was observed upon the depletion of Sdo1, an essential cofactor for ribosomal activation^58,59^, or Lsg1, a GTPase required to license the subunit for 40S joining^60,61^, using an auxin-inducible degron (AID) system^62^ (Figure 5E, lanes 1 and 4; verification of the depletion in Figure S7B). These results establish that maturation stalls at multiple distinct steps of the pre-60S pathway serve as a general trigger for Reh1-Mms1-mediated ubiquitination, firmly positioning Reh1 as a universal beacon for defective 60S subunits.

Collectively, these findings converge on a unified model for ribosomal quality control (Figure 6). We propose that Reh1 acts as a dynamic sensor that monitors the timely progression of both ribosome maturation and nascent chain synthesis. Under normal conditions, the successful completion of maturation and the subsequent production of a nascent chain displace Reh1 from the PET, ensuring that the E3 machinery is not recruited. However, when the process is interrupted—either by assembly failures that stall pre-60S maturation or by nonfunctional 25S rRNAs that impair nascent chain production—the prolonged occupancy of Reh1 provides a critical time window for the recruitment of the Mms1 (E3) complex. This “surveillance-by-retention” mechanism allows the cell to safeguard the integrity of the protein synthesis machinery by clearing ribosomes that fail to reach functional maturity.

**Figure 6.**
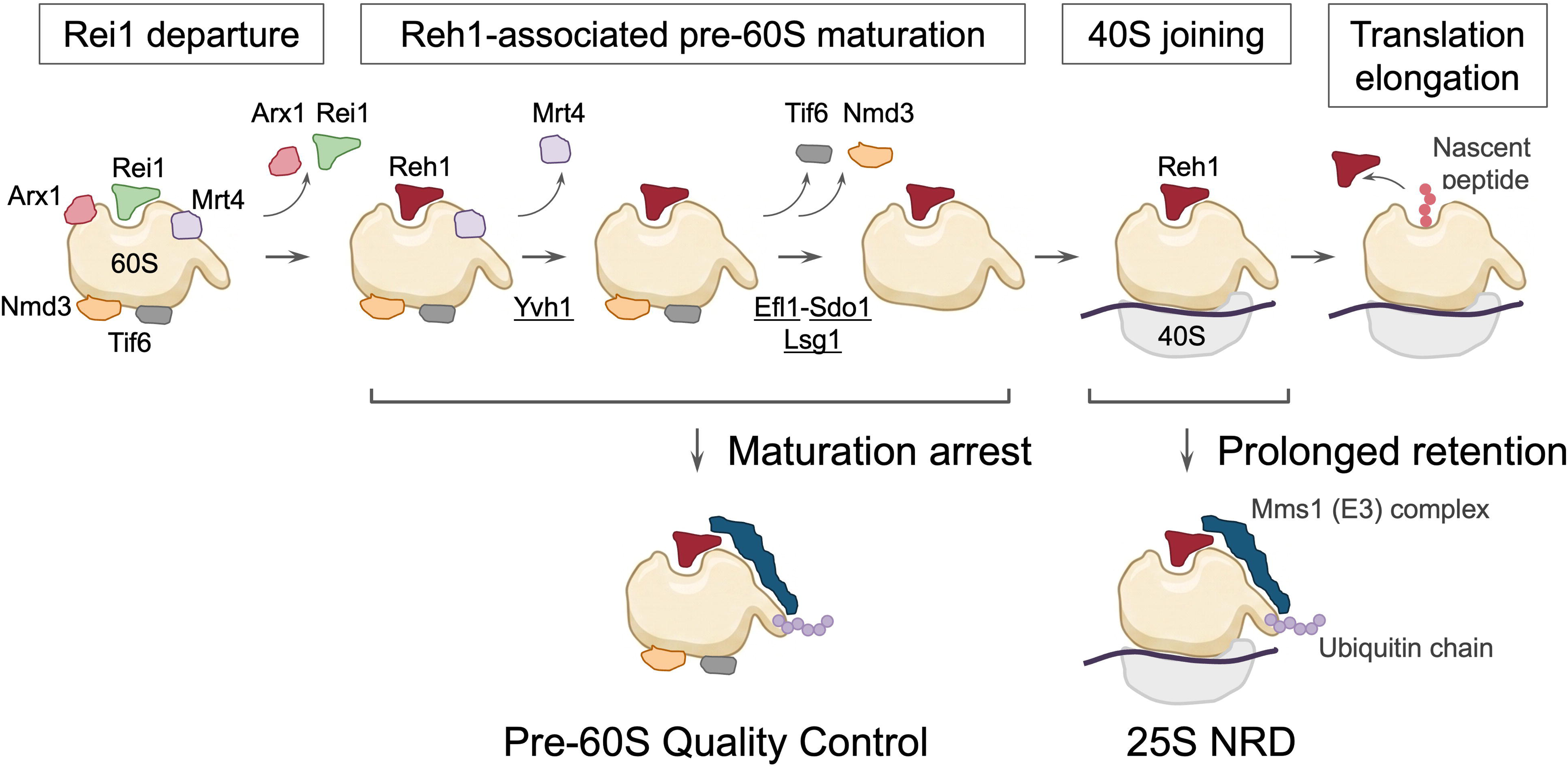
A unified “surveillance-by-retention” model for 60S quality control. Schematic of the proposed model. Reh1 acts as a universal timer by monitoring the polypeptide exit tunnel (PET). Successful maturation and translation initiation lead to Reh1 eviction by the nascent chain. Conversely, maturation stalls (e.g., in *yvh1*Δ, *efl1*, or *sdo1/lsg1* mutants) or translational incompetence (nonfunctional 25S rRNA) cause prolonged Reh1 retention, providing a time window for Mms1 recruitment, Rpl19 ubiquitination, and subsequent subunit degradation.

## Discussion

In this study, we have unraveled the molecular logic of 25S nonfunctional rRNA decay (25S NRD) and identified the ribosome biogenesis factor Reh1 as a specialized sensor for defective large subunits. This crucial role of Reh1 as an NRD factor is corroborated by a recent bioRxiv paper showing that the *reh1*Δ strain extends the half-life of 25S NRD substrates^63^. We demonstrated that Reh1 physically bridges the Mms1-Rtt101-Crt10 complex to nonfunctional 80S ribosomes via its peptide exit tunnel (PET) insertion, directly guiding the specific ubiquitination of Rpl19 at its C-terminal extension to mark the particle for degradation. Furthermore, we discovered that this quality control response is unexpectedly triggered during delays in late pre-60S maturation stages, thereby extending the significance of this pathway beyond NRD to a broader surveillance of ribosome assembly.

The selective protection of functional 80S particles from ubiquitination is rationalized by the programmed departure of Reh1, driven by the synthesis of the nascent peptide chain. Our findings support a model where the elongating nascent peptide directly competes with the C-terminal helix of Reh1 within the PET (Figure 6). This spatial competition ensures that Reh1 is efficiently displaced from functional 80S ribosomes but remains exclusively retained on nonfunctional ones (Figure 3D)—such as those stalled at the start codon with an empty PET—thereby triggering the Mms1-dependent decay pathway.

Structurally mature but catalytically inactive 60S subunits readily assemble with functional 40S subunits to form 80S complexes at the initiation codon. Upon arrival of the second aminoacyl-tRNA, the wild-type decoding center hosted by the 40S subunit accepts it normally, triggering the corresponding conformational rearrangements^64–66^. Only after this step does the functional defect of the PTC manifest, trapping the ribosome in a stalled state with the aminoacyl-tRNA fully inserted into the A-site. Because this state lacks a vacant A-site, it remains completely invisible to sensors like GCN1^18^. This structural bottleneck highlights the supreme rationality of the PET-monitoring mechanism discovered here, which uniquely leverages tunnel vacancy to flag large subunit dysfunction.

Another key discovery is that the Reh1-Mms1 pathway serves as a broad-spectrum surveillance system for pre-60S biogenesis, as evidenced by the massive, Reh1-dependent ubiquitination triggered by various late-stage maturation stalls (e.g., in yvh1Δ, lsg1-aid, or sdo1-aid cells; Figure 5). This raises a fundamental question: how does the E3 complex discriminate between stalled particles and normal maturation intermediates, given that Reh1 transiently associates with both? Our findings suggest that this selectivity is governed by a kinetic “time-window” rather than a specific inhibitory mask (Figure 6). Under normal conditions, the residence time of Reh1 on a maturing particle is likely too brief for the stochastic recruitment of the Mms1 complex. However, any maturation stall—whether caused by a factor deficiency or a structural defect—extends this occupancy, providing the critical time required for the low-abundance Mms1 complex to encounter and bind to Reh1’s N-terminal domain, thereby initiating the E3-mediated response. This “surveillance-by-retention” mechanism thus allows the cell to effectively distinguish between productive assembly and terminal stagnation. This concept aligns with emerging paradigms in higher eukaryotes, such as the mammalian RASP pathway where dead-end biogenesis intermediates are actively targeted for ubiquitin-dependent degradation^67^, emphasizing that sorting flawed assembly from productive maturation is a universally critical task.

The specific ubiquitination of Rpl19 at its extreme C-terminus provides profound mechanistic insight into the downstream fate of the defective ribosome. The targeted lysines (K180 and K186) are located precisely at the eB12 inter-subunit bridge (Figure 4E), where Rpl19 physically contacts the 40S subunit^4,68^. We previously demonstrated that the Cdc48-Ufd1-Npl4 segregase complex is strictly required to dissociate non-functional 80S ribosomes into distinct subunits prior to proteasomal degradation^28^. The spatial arrangement discovered here is highly suggestive: by specifically ubiquitinating the eB12 bridge, the Mms1 complex places the degradation signal exactly at the subunit interface. We propose that this site-specific ubiquitination directly recruits the Cdc48 segregase to the precise location where mechanical force is needed to wedge apart the stalled, defective 80S particle.

The mechanistic elegance of this system is highlighted by the functional divergence between Reh1 and its paralog Rei1. Both proteins utilize a conserved C-terminal tail to probe the PET, yet they monitor the tunnel with opposite biological objectives: Rei1 licenses structural maturation by sensing the “openness” of the tunnel to trigger Arx1 release^45,46^, whereas Reh1 subsequently verifies functional competence by detecting the “idleness” of the same tunnel. This transition—from a factor that promotes biogenesis to one that validates operation—represents a sophisticated evolutionary partition of PET-sensing duties. The physiological importance of this specialization is underscored by the distinct regulatory circuits of these two paralogs; while *REI1* is co-regulated with constitutive biogenesis factors, transcriptomic data reveal that *REH1* expression is uniquely synchronized with the ubiquitin-proteasome system (UPS), establishing it as a dedicated quality control factor.

Our study thus reveals a surveillance system where a “retained” maturation factor serves as a universal mark for ribosomal failure. Each pre-60S particle receives Reh1 as a potential degradation trigger—a fail-safe strategy that ensures no dysfunctional particle leaks into the translating pool. Such a stringent system highlights the critical importance of ribosomal integrity, as evidenced by the severe human ribosomopathies^69,70^ caused by even subtle defects.

Furthermore, this mechanism of “rsurveillance-by-retention” may represent a broader paradigm for the quality control of other large macromolecular complexes that require high-fidelity assembly and function.

## Limitations of the Study

Several limitations of this study should be noted. First, our findings are based on the budding yeast Saccharomyces cerevisiae; further research is needed to determine if this “surveillance-by-retention” mechanism is identical in higher eukaryotes. Second, while we identified the Rpl19 ubiquitination site favorable for Cdc48 recognition, the exact functional roles of Cdc48 and the downstream ribonucleases clearing the defective 25S rRNA remain to be characterized. Finally, although CRAC mapping successfully resolved the spatial proximity between the Reh1 N-terminus and Rpl19, obtaining a high-resolution atomic model of this dynamic complex via cryo-EM remains a future challenge due to its high conformational flexibility.

## Acknowledgements

We thank Drs. Ichiro Taniguchi (Osaka University) and Mutsuhito Ohno (Kyoto University) for helpful discussions, and Dr. Rina Hamajima (Nagoya University) for technical assistance and discussions. We also thank Dr. Akira Nakamura and Ms. Kaori Shinmyozu (Riken, Kobe) for their assistance with the MS analysis. The BYP6998 plasmid and the BY25598 yeast strain were provided by the National Bio-Resource Project (NBRP), Japan. This research was supported in part by JSPS KAKENHI (grant numbers 24657117, 26440004, 19K06487, 22K06080 and 26K09208 to M.K.); by Grants-in-Aid for Scientific Research on Innovative Areas from MEXT, Japan (grant numbers 21112511, 23112710); by the Takeda Science Foundation to M.K.; and by PRESTO, Japan Science and Technology Agency (JST), to M.K.

## Author contributions

T.S. performed the key experiments. M.K. performed the remaining experiments and analyzed the data. K.F. contributed to data analysis, project management, and interpretation of results. M.K. drafted the manuscript, and K.F. critically revised it.

## STAR⍰METHODS

### KEY RESOURCES TABLE

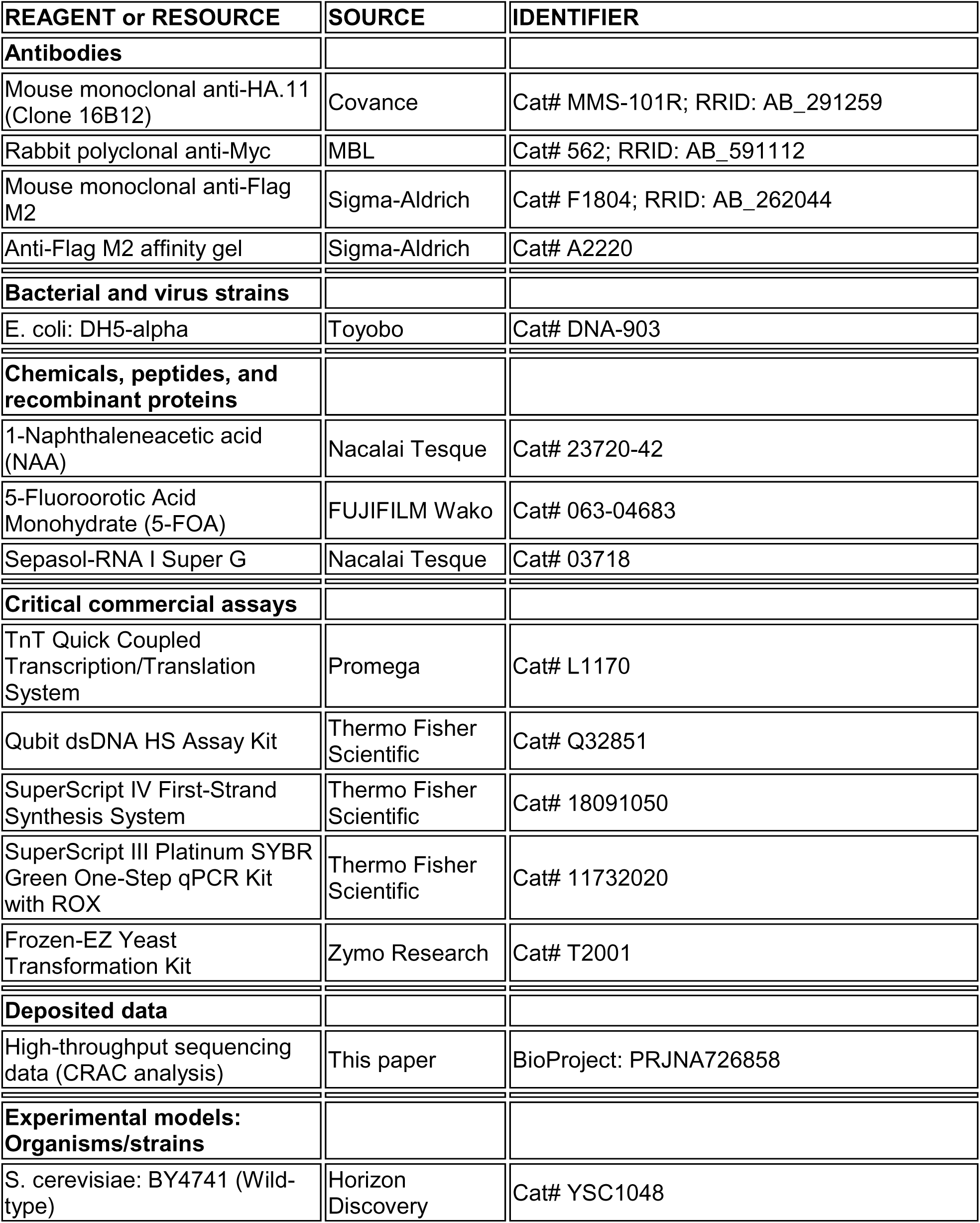
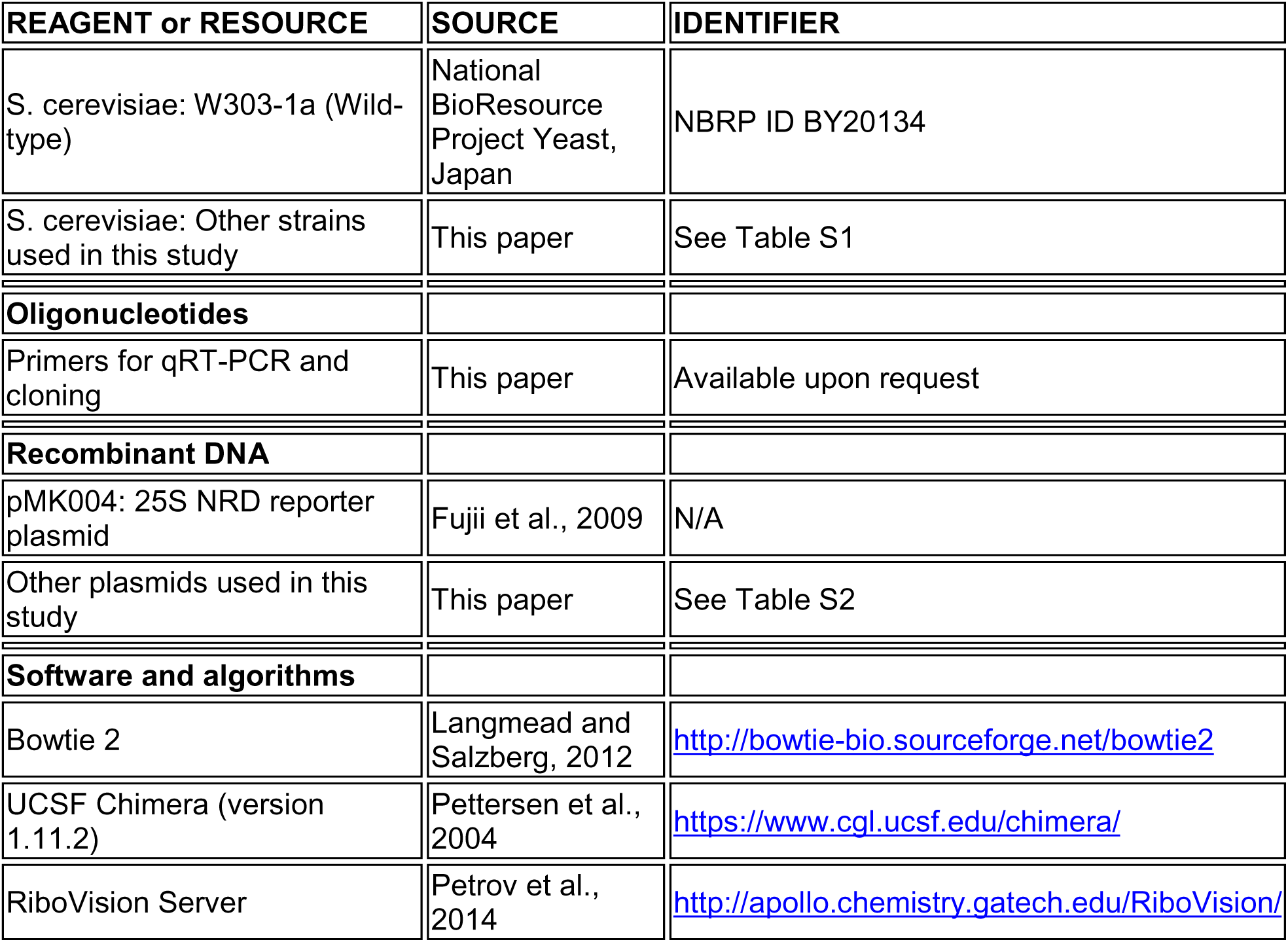

| REAGENT or RESOURCE | SOURCE | IDENTIFIER |
| --- | --- | --- |
| **Antibodies** |  |  |
| Mouse monoclonal anti-HA.11 (Clone 16B12) | Covance | Cat# MMS-101R; RRID: AB_291259 |
| Rabbit polyclonal anti-Myc | MBL | Cat# 562; RRID: AB_591112 |
| Mouse monoclonal anti-Flag M2 | Sigma-Aldrich | Cat# F1804; RRID: AB_262044 |
| Anti-Flag M2 affinity gel | Sigma-Aldrich | Cat# A2220 |
| **Bacterial and virus strains** |  |  |
| E. coli: DH5-alpha | Toyobo | Cat# DNA-903 |
| **Chemicals, peptides, and recombinant proteins** |  |  |
| 1-Naphthaleneacetic acid (NAA) | Nacalai Tesque | Cat# 23720-42 |
| 5-Fluoroorotic Acid Monohydrate (5-FOA) | FUJIFILM Wako | Cat# 063-04683 |
| Sepasol-RNA I Super G | Nacalai Tesque | Cat# 03718 |
| **Critical commercial assays** |  |  |
| TnT Quick Coupled Transcription/Translation System | Promega | Cat# L1170 |
| Qubit dsDNA HS Assay Kit | Thermo Fisher Scientific | Cat# Q32851 |
| SuperScript IV First-Strand Synthesis System | Thermo Fisher Scientific | Cat# 18091050 |
| SuperScript III Platinum SYBR Green One-Step qPCR Kit with ROX | Thermo Fisher Scientific | Cat# 11732020 |
| Frozen-EZ Yeast Transformation Kit | Zymo Research | Cat# T2001 |
| **Deposited data** |  |  |
| High-throughput sequencing data (CRAC analysis) | This paper | BioProject: PRJNA726858 |
| **Experimental models: Organisms/strains** |  |  |
| S. cerevisiae: BY4741 (Wild-type) | Horizon Discovery | Cat# YSC1048 |
| *S. cerevisiae*: W303-1a (Wild-type) | National BioResource Project Yeast, Japan | NBRP ID BY20134 |
| *S. cerevisiae*: Other strains used in this study | This paper | See Table S1 |
| **Oligonucleotides** |  |  |
| Primers for qRT-PCR and cloning | This paper | Available upon request |
| **Recombinant DNA** |  |  |
| pMK004: 25S NRD reporter plasmid | Fujii et al., 2009 | N/A |
| Other plasmids used in this study | This paper | See Table S2 |
| **Software and algorithms** |  |  |
| Bowtie 2 | Langmead and Salzberg, 2012 | http://bowtie-bio.sourceforge.net/bowtie2 |
| UCSF Chimera (version 1.11.2) | Pettersen et al., 2004 | https://www.cgl.ucsf.edu/chimera/ |
| RiboVision Server | Petrov et al., 2014 | http://apollo.chemistry.gatech.edu/RiboVision/ |

### RESOURCE AVAILABILITY

#### Lead Contact

Further information and requests for resources and reagents should be directed to and will be fulfilled by the lead contact, Makoto Kitabatake.

#### Materials Availability

All unique/stable reagents generated in this study are available from the lead contact without restriction.

#### Data and Code Availability

- High-throughput sequencing data (CRAC analysis) have been deposited at the NCBI Sequence Read Archive (SRA) under BioProject: PRJNA726858 and are publicly available as of the date of publication.
- This paper does not report original code.
- Any additional information required to reanalyze the data reported in this paper is available from the lead contact upon request.

### EXPERIMENTAL MODEL AND SUBJECT DETAILS

#### Yeast Strains (Saccharomyces cerevisiae)

All Saccharomyces cerevisiae strains used in this study are listed in Table S1. Strains NOY401 (utilized for the RNA polymerase I temperature-sensitive rescue assay (Figure 1B and Figures S1C and S1D)) and BY25598 (utilized for auxin-inducible degron (AID) experiments (Figure 5E and Figure S7B)), along with all strains constructed within these genetic backgrounds (MKY23, MKY458, MKY461, MKY462, MKY489, and MKY490), were derived from W303-1a (MATa ade2-1 his3-11,15 leu2-3 trp1-1 ura3-1 can1-100). Other strains were generated from the BY4741 background (MATa his3Δ1 leu2Δ0 met15Δ0 ura3Δ0). Genomic deletions or epitope tag modifications were performed via standard PCR-mediated homologous recombination. Transformants were selected using appropriate drug-resistance or auxotrophic markers, and successful genomic integration was verified by colony PCR. Strains obtained from the Yeast Knock-out MATa Collection (Horizon Discovery, Cambridge, UK) were validated using the same verification process. Due to an inability to confirm the designated genomic sequence of the corresponding strain from our initial stock, a slm6Δ strain was utilized as a rei1Δ strain after genetic verification (Figure 2C and Figure S3C, S3D). In this slm6Δ strain, the SLM6 gene is encoded on the complementary strand of the REI1 locus, and a 453-bp fragment within the 5′ region of the 1,182-bp REI1 open reading frame (ORF) was replaced by the selection cassette.

#### Bacterial Strains (Escherichia coli)

Escherichia coli strain DH5-alpha was used for plasmid construction and subsequent plasmid propagation. Cells were cultured in LB liquid medium or on LB agar plates supplemented with appropriate antibiotics at 37°C, unless otherwise specified.

### METHOD DETAILS

#### Yeast Culture Conditions and Media

Yeast strains were grown in rich medium (YPD; 1% yeast extract, 2% peptone, and 2% glucose) or synthetic drop-out (SD) medium containing 2% glucose and the appropriate amino acid supplements. Transformations were performed using the Frozen-EZ Yeast Transformation Kit (Zymo Research, Irvine, CA, USA). For galactose-inducible expression driven by the GAL1 or GAL7 promoters, colonies from an SD-glucose plate were pre-cultured overnight in SD-raffinose (2%) medium. Expression was induced by inoculating pre-cultures into large-scale SD-galactose (2%) medium at an initial OD600 of 0.05. Cultures were harvested during the mid-logarithmic growth phase (OD600 = 0.5–0.7). For AID-mediated depletion experiments (Figure 5E and Figure S7B), pre-cultures grown in SD-glucose were inoculated into fresh SD-glucose medium supplemented with 0.05 mM 1-Naphthaleneacetic acid (NAA; Nacalai Tesque, Kyoto, Japan) at an initial OD600 of 0.05 and grown to mid-log phase. Unless otherwise specified, all liquid cultures were incubated at 30°C.

#### Colony Formation Assays

Rescue experiments for the RNA polymerase I temperature-sensitive strain (Figure 1B and Figure S1A) were performed on SD-galactose plates. Phenotypic analysis of the reh1Δ, rei1Δ, and yvh1Δ strains (Figure S3D) and the rei1Δ rescue experiments (Figure S3C) were conducted on YPD plates. Other phenotypic assays (such as plasmid shuffling counterscreening) were carried out on the indicated selective media (Figure S5B). Overnight cultures were serially diluted 10-fold with sterile water, and 2.5 μL of each dilution was spotted onto plates. Plates were incubated at 30°C (unless otherwise indicated) for 2 days prior to digital imaging.

#### Plasmid Construction and Shuffling

Plasmids used in this study were constructed using standard molecular cloning protocols, and all insert sequences were verified by Sanger sequencing. A comprehensive list of plasmids is provided in Table S2. The structural organization of the 25S NRD reporter insert (pMK004) is illustrated in Figure 1A. The sequence and precise position of the 18-nt tag sequence within the 25S rRNA, along with the sequences of northern blotting probes and qRT-PCR primers, were previously described (Fujii et al., 2009). Mutations surrounding the peptidyl transferase center (PTC) were introduced via overlap extension PCR.

For low-copy expression, the ORFs of interest, along with approximately 500 bp of upstream and downstream flanking genomic regions for *REH1*, *REI1*, *YVH1*, *RPL11A*, *MRT4*, *LSG1*, *SDO1*, and *RPL19A*, were amplified by PCR and cloned into CEN-type vectors (pMK064 and pMK069 backgrounds). Epitope tags were introduced at either the N- or C-terminus of the ORF as required. For the *RPL19A* expression plasmids (pMK045 and its derivatives), the endogenous downstream flanking region was replaced with that of *RPL13A* to facilitate cloning.

For high-copy overexpression, the *GAL1* or *GAL7* promoter was inserted upstream of the target gene. The Mms1-Flag expression plasmid (pMK364) was generated by replacing the 3′ TAP-tag sequence of a commercial Mms1-TAP plasmid (BG1805 derivative, Yeast ORF Collection, Horizon Discovery) with a Flag-tag sequence. The Rtt101 expression vector (pMK365) was constructed from a commercial Rtt101-TAP plasmid by homologous recombination with a PCR fragment amplifying the untagged, endogenous *RTT101* gene. The Crt10-HA expression plasmid (pMK363) was constructed using a similar strategy. For overexpression of the GST-tagged Efl1 protein, a fragment encompassing the GST ORF and the multiple cloning site from pGEX-6P-1 (Cytiva, Tokyo, Japan) was cloned into the HpaI-HindIII sites of YEplac195, yielding pMK351. The *EFL1* ORF was amplified from genomic DNA and inserted into the SalI-NotI sites of pMK351 to generate pMK354 (Figure 5D and Figure S7A). All primer sequences are available upon request.

For plasmid shuffling experiments (Figures S5A and S5B), the wild-type *RPL19A* gene was cloned into the *URA3*-marked CEN vector pRS316 and introduced into an *rpl19a*Δ strain prior to the genomic disruption of *RPL19B*. Successful deletion of *RPL19B* was verified by PCR. Next, *rpl19a* mutant alleles cloned into the *LEU2*-marked CEN vector YCplac111 were transformed into the resulting *rpl19a*Δ *rpl19b*Δ double-deletion strain harboring pRS316-RPL19A. To allow the spontaneous loss of the *URA3* plasmid, cells were cultured in medium selective only for the *LEU2* plasmid (SD–Leu). Serial dilutions of the cells were then spotted onto synthetic complete (SC) agar plates containing 0.1% 5-fluoroorotic acid (5-FOA) and 2% glucose to counter-select against the wild-type *URA3* plasmid.

#### Isolation of Active E3 Ubiquitin Ligase Complex

To isolate the active E3 ubiquitin ligase complex (Mms1-Flag, Rtt101, and Crt10-HA) (Figures 2A and 2B), the respective expression vectors were co-transformed into a *paf1*Δ strain, as the E3 complex associates with the Paf1 complex *in vivo*, which is dispensable for 25S NRD (Sakata et al., 2015). A strain lacking the Mms1-Flag plasmid served as a negative control.

Strains were pre-cultured in SD-raffinose and inoculated into SD-galactose medium to induce the *GAL1* promoter. Cells were grown at 30°C to an OD600 of 0.8, harvested, and ground into a fine powder in a mortar under liquid nitrogen. Total proteins were extracted using IPP150 buffer [10 mM Tris-HCl (pH 7.5), 2.5 mM MgCl2, 150 mM NaCl, 0.1% Nonidet P-40]. The active E3 complex was isolated using anti-Flag M2 affinity gel (Sigma-Aldrich, St. Louis, MO, USA) and washed extensively with IPP150 buffer before being subjected to *in vitro* binding assays.

#### Cell-Free Protein Synthesis and *In Vitro* Binding Assays

Radiolabeled 35S-labeled proteins (candidate factors and Reh1 truncation mutants) were synthesized using the TnT Quick Coupled Transcription/Translation T7 System (Promega, Madison, WI, USA) with PCR-amplified DNA templates. Following a 90-min translation reaction at 30°C, samples were diluted in IPP150 buffer and incubated with the active Mms1-Flag/Rtt101/Crt10-HA E3 complex immobilized on anti-Flag M2 agarose beads (Figures 2A and 2B). Mixtures were rotated gently at 4°C for 60 min, and the beads were harvested and washed extensively with either standard IPP150 buffer or IPP150 containing 500 mM NaCl to remove non-specific interactors. Bound proteins were resolved by 12.5% SDS-PAGE and visualized by autoradiography. Images were acquired using a Typhoon FLA 7000 imager (Cytiva) at multiple exposure times, and panels displaying equivalent input intensities were selected for presentation (Figures 2A and 2B).

#### Northern Hybridization and Quantitative RT-PCR

Total RNA was extracted from yeast cultures harvested at mid-log phase using the proteinase K method and quantified via NanoDrop (Thermo Fisher Scientific, Waltham, MA, USA). Northern hybridization and quantitative RT-PCR (qRT-PCR) analyses were performed as previously described (Fujii et al., 2009). For northern blotting (Figure 1B, Figure 4D, and Figure S1A), sample loading was normalized to the culture OD600 at the time of harvest (400 ng of total RNA for the t = 0 time point). RNA was resolved on a 1% agarose gel, transferred to a Hybond-N+ membrane (Cytiva) via capillary transfer, and probed for the 18-nt tag sequence of the 25S NRD reporter. For qRT-PCR (Figure 2C, Figures 3B–3D, and Figures S2A and S2B), total RNA was analyzed using two primer sets targeting either the 18-nt tagged or untagged endogenous 25S rRNA. Assays were conducted on a StepOnePlus Real-Time PCR System using the SuperScript III Platinum SYBR Green One-Step qPCR Kit with ROX (Thermo Fisher Scientific). All qRT-PCR experiments were performed in at least biological triplicate using independent colonies. Tagged rRNA levels were normalized to untagged endogenous 25S rRNA to control for extraction efficiency. To account for variations in *GAL7* promoter induction and growth rates among different mutant strains, relative mutant 25S rRNA stability was determined by calculating the ratio of mutant to wild-type tagged 25S rRNA (Figure 2C and Figure S2). For analysis of immunoprecipitated complexes, bound RNA was eluted from beads and purified using Sepasol-RNA I Super G (Nacalai Tesque). Northern blotting of sucrose density gradient fractions was conducted as previously described (Fujii et al., 2009).

#### Protein Extraction, Immunoblotting, and Immunoprecipitation

Proteins were resolved by 5%–20% SDS-PAGE and transferred to 0.2-μm Amersham Protran nitrocellulose membranes (Cytiva) using a semi-dry transfer method. Membranes were probed with the following primary antibodies: anti-HA.11 monoclonal antibody (16B12; Covance, Princeton, NJ, USA; used for detecting Rpl19-HA variants and candidate factors (Figures 4A and 4B, and Figure S4D)); anti-Myc polyclonal antibody (rabbit; MBL, code no. 562; used for total ribosomal ubiquitination (Figure 2D, Figure 4C, Figures 5B–5E, and Figures S4A and S4B)); and anti-Flag M2 monoclonal antibody (F1804; Sigma-Aldrich; used for confirming ribosome purification and E3 isolation). For anti-Flag immunoprecipitations, lysates were incubated with anti-Flag M2 affinity gel (A2220; Sigma-Aldrich) at 4°C, and the immune complexes were washed three times with IPP150 buffer prior to elution.

#### Sucrose Density Gradient Centrifugation and Ribosome Isolation

Ribosomal 60S and 80S complexes were isolated from clarified cell lysates by 10%–40% sucrose density gradient ultracentrifugation using an SW41 rotor (Beckman Coulter) at 273,620 × g for 150 min at 4°C (Figures 3A, 3C, 3D, 5C, 5D, 5E, S3D, and S7A). Gradients were fractionated into 85 fractions using a BioComp Gradient Station (Fredericton, NB, Canada) while monitoring absorbance at 260 nm. Fractions corresponding to 60S or 80S peaks were pooled, concentrated using Amicon Ultra 0.5-mL 100K centrifugal filters (Merck, Darmstadt, Germany), and quantified via NanoDrop. Ribosomal particles were adjusted to a final concentration of 50 nM in detergent-free IPP150 buffer, and 7-μL aliquots per lane were analyzed by immunoblotting.

#### Identification of Ubiquitination Targets by Mass Spectrometry

The BY4741 wild-type strain was co-transformed with pMK091 (encoding His6-Myc-ubiquitin G76A under the control of the CUP1 promoter) and pMK272 (the 25S NRD reporter) and cultured overnight in SD-raffinose. To overproduce the mutant 25S rRNA, pre-cultures were inoculated into a total volume of 24 L of SD-galactose medium at an initial OD600 of 0.05 and grown at 30°C to an OD600 of ∼0.5. Expression of His6-Myc-ubiquitin was subsequently induced by adding CuSO4 to a final concentration of 0.1 mM, and cells were cultured for an additional 3 h prior to harvesting. Frozen cell pellets were pulverized in a mortar under liquid nitrogen, and total proteins were extracted with 24 mL of IPP150 buffer. The 60S ribosomal fraction was isolated via 10%–40% sucrose gradient ultracentrifugation using an SW28 rotor (Beckman Coulter) at 131,101 × g for 5 h at 4°C in the presence of 40 mM EDTA. The 60S fractions were pooled, precipitated with trichloroacetic acid (TCA; 15% final concentration), washed with ice-cold acetone, and resuspended in IPP150 buffer containing 8 M urea.

Ubiquitinated proteins were purified by HisTrap column chromatography (Cytiva) using a 10–500 mM linear imidazole gradient (Figures S4A and S4B). Samples were processed using the Pierce SDS-PAGE Sample Prep Kit (Thermo Fisher Scientific), resolved by 5%–20% SDS-PAGE, and visualized by silver staining (Figure S4B). Protein bands unique to the target fractions were excised and analyzed by liquid chromatography-tandem mass spectrometry (LC-MS/MS). MS/MS data were processed and searched against the NCBInr database (release 20100403; restricted to the Saccharomyces cerevisiae taxonomy, 26,164 sequences) using Mascot Daemon (Matrix Science). The search parameters were set as follows: trypsin digestion with a maximum of one missed cleavage; fixed modification of carbamidomethylation on cysteine; variable modification of oxidation on methionine; peptide mass tolerance of ± 2 Da; and fragment mass tolerance of ± 0.8 Da using monoisotopic mass values. For each of the 11 candidate proteins identified by MS analysis (Figure S4C), a C-terminally HA-tagged expression vector was constructed for downstream individual biochemical validation (Figure 4A and Figure S4D).

#### UV Cross-linking and cDNA Analysis (CRAC)

In vivo CRAC experiments were performed as previously described (Granneman et al., 2009) with modifications. The reh1Δ or rei1Δ strains were transformed with pMK452 (encoding Flag-His6-Reh1) or pMK451 (encoding Flag-His6-Rei1), respectively; an untransformed BY4741 strain served as a negative control. Cells grown to mid-log phase were irradiated in vivo with 1.6 J/cm2 of UVC (254 nm) using an FS-800 UV Crosslinker (Funakoshi, Tokyo, Japan) and snap-frozen in liquid nitrogen. For each sample, 1 g of frozen cell pellet was pulverized under liquid nitrogen and extracted with IPP150 buffer. Lysates were subjected to immunoprecipitation using 80 μL of anti-Flag M2 agarose beads. Beads were washed with IPP150 buffer, equilibrated in 1× PNK buffer, and RNAs were partially digested with RNace-It Ribonuclease Cocktail (3 U; Agilent Technologies) for 5 min at 37°C with agitation. The reaction was quenched by adding 9 volumes of 1× wash buffer containing 6 M guanidine-HCl, and eluates were filtered. Eluates were incubated with Ni-NTA agarose (Qiagen, Hilden, Germany) at 4°C and processed as described (Granneman et al., 2009), except that T4 polynucleotide kinase (Toyobo, Osaka, Japan) was applied to dephosphorylate the 3′ ends of short RNA tags. A 5′ DNA Adenylation Kit (New England Biolabs, Ipswich, MA, USA) was used to prepare the 5′ adenylated 3′-linker. cDNA synthesis was performed using the SuperScript IV First-Strand Synthesis System (Thermo Fisher Scientific), and cDNA libraries were amplified by PCR using EmeraldAmp DNA Polymerase (Takara Bio, Kyoto, Japan). The resulting 150–175 bp PCR products were purified by 8% PAGE and selected using SPRIselect beads (1.2×; Beckman Coulter) after a second round of amplification with PrimeSTAR GXL (Takara Bio) using minimal cycle numbers to maintain non-saturating conditions. Final libraries were quantified using the Qubit DNA HS Assay Kit on a Qubit 3 Fluorometer (Thermo Fisher Scientific), pooled in equimolar ratios, and sequenced on an Illumina NovaSeq platform (Gene Nex, Kumamoto, Japan) using a paired-end 150-bp (PE150) configuration, from which single-end 50-bp (SE50) equivalent datasets were extracted and analyzed. After demultiplexing, reads containing both 5′ and 3′ adapters (8–32 nt inserts) were selected. Reads were mapped to the Saccharomyces cerevisiae S288C reference genome (NCBI build 3.1) using Bowtie2, allowing one mismatch per read. Resolution of the 35S rDNA repeat unit is shown in Figure S6A. Consensus sequences are mapped onto the rRNA secondary structure in Figure S6B. Two independent biological replicates encompassing independent cultures, UV irradiation, and sequencing were performed and yielded identical results. The rRNA secondary structure maps were generated using the RiboVision Server (Petrov et al., 2014). Three-dimensional structural models of the ribosome (Figure 4E and Figure S6C) were rendered in UCSF Chimera (version 1.11.2) using coordinates from PDB: 4V88; the 40S subunit and Stm1 were omitted for clarity.

#### Bioinformatics and Co-expression Analysis

Co-expression analysis of REI1 and REH1 was performed using the SPELL (Serial Pattern of Expression Levels Locator) tool on the SGD platform (Hibbs et al., 2007; Wong et al., 2023). REI1 and REH1 were entered as query genes to evaluate their expression correlation across available microarray and RNA-seq datasets.

### QUANTIFICATION AND STATISTICAL ANALYSIS

#### Data Reporting

No statistical methods were used to predetermine sample sizes. The experiments were not randomized, and the investigators were not blinded to allocation during experiments and outcome assessment. Quantitative qRT-PCR assays were performed in at least biological triplicate starting from independent yeast colonies, and data are presented as mean ± SD where applicable, as indicated in the respective figure legends. Statistical significance between specific groups was evaluated where appropriate.

