## Supplementary Material for "Absence of nascent peptides triggers nonfunctional ribosome decay"

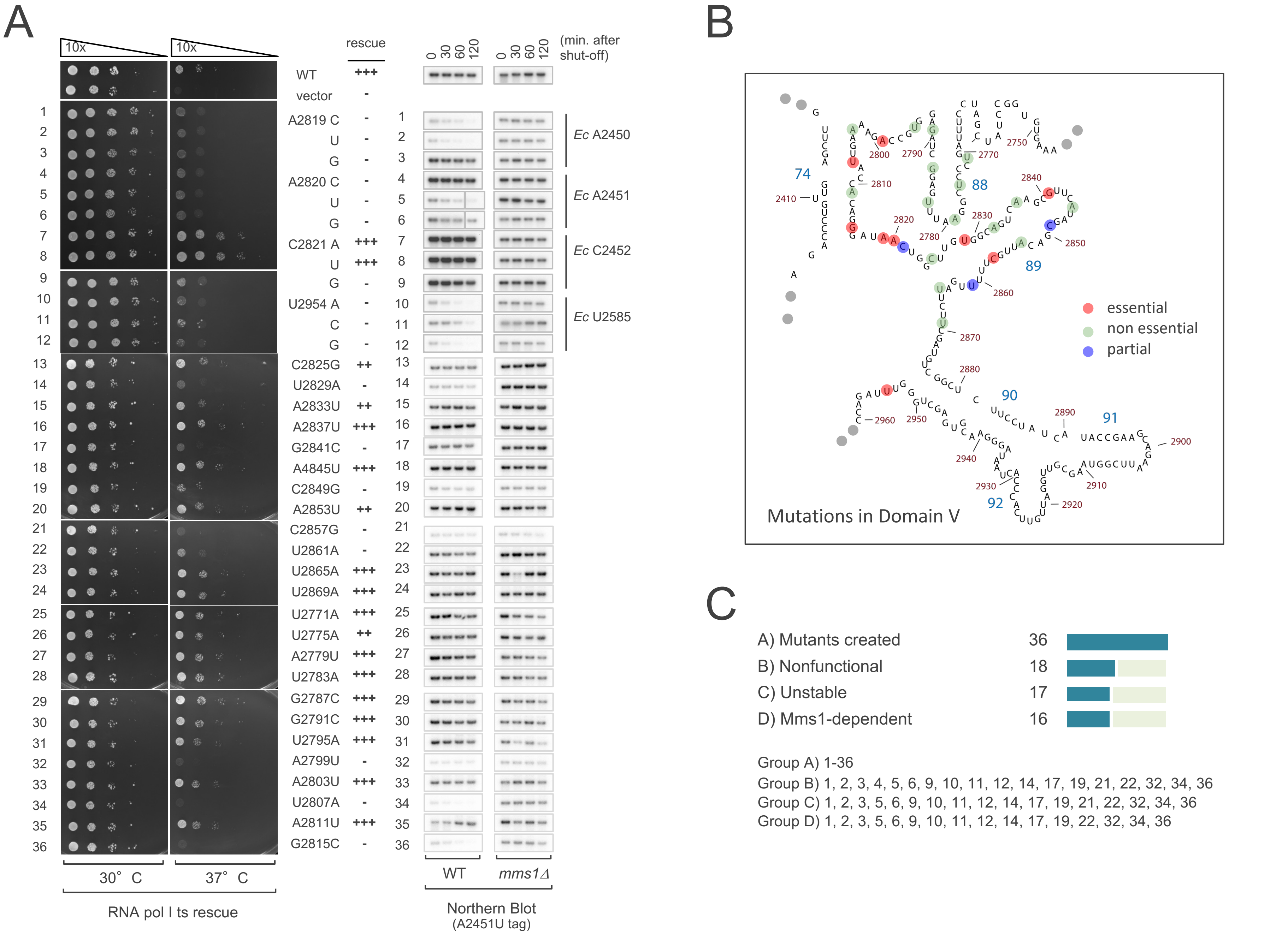
A
10x
10x
rescue
0 30 60 120
0 30 60 120
(min. after shut-off)
WT +++
vector -
A2819 C -
U -
G -
A2820 C -
U -
G -
C2821 A +++
U +++
G -
U2954 A -
C -
G -
C2825G ++
U2829A -
A2833U ++
A2837U +++
G2841C -
A4845U +++
C2849G -
A2853U ++
C2857G -
U2861A -
U2865A +++
U2869A +++
U2771A +++
U2775A ++
A2779U +++
U2783A +++
G2787C +++
G2791C +++
U2795A +++
A2799U -
A2803U +++
U2807A -
A2811U +++
G2815C -
Ec A2450
Ec A2451
Ec C2452
Ec U2585
30° C
37° C
RNA pol I ts rescue
WT
mms1Δ
Northern Blot
(A2451U tag)
B
74
88
89
90
91
92
essential
non essential
partial
Mutations in Domain V
C
A) Mutants created 36
B) Nonfunctional 18
C) Unstable 17
D) Mms1-dependent 16
Group A) 1-36
Group B) 1, 2, 3, 4, 5, 6, 9, 10, 11, 12, 14, 17, 19, 21, 22, 32, 34, 36
Group C) 1, 2, 3, 5, 6, 9, 10, 11, 12, 14, 17, 19, 21, 22, 32, 34, 36
Group D) 1, 2, 3, 5, 6, 9, 10, 11, 12, 14, 17, 19, 22, 32, 34, 36

Figure S1

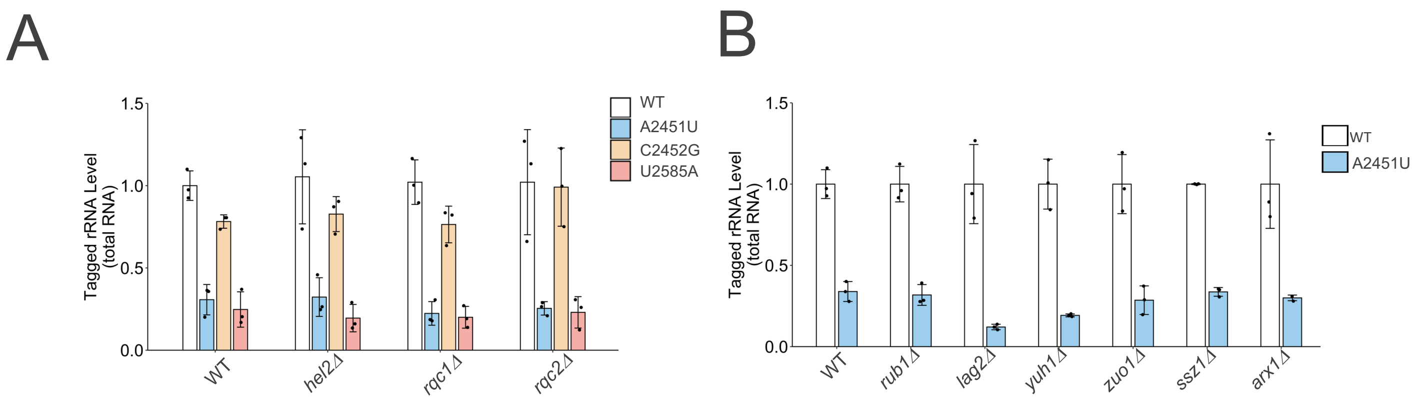
A
B
Tagged rRNA Level (total RNA)
1.5
1.0
0.5
0.0
WT
A2451U
C2452G
U2585A
WT
*hel2Δ*
*rqc1Δ*
*rqc2Δ*
Tagged rRNA Level (total RNA)
1.5
1.0
0.5
0.0
WT
A2451U
WT
*rub1Δ*
*lag2Δ*
*yuh1Δ*
*zuo1Δ*
*ssz1Δ*
*arx1Δ*

Figure S2

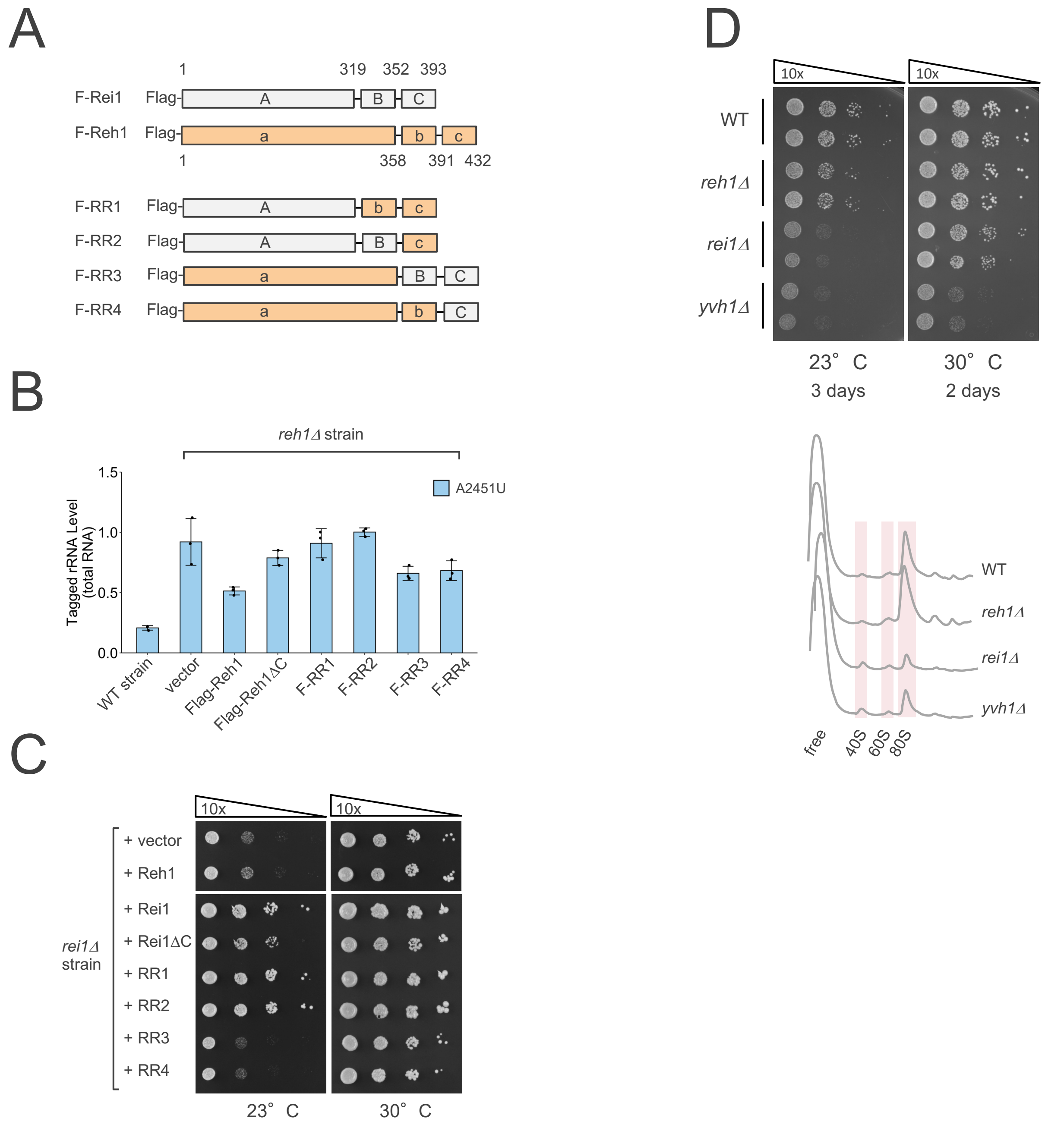
A
1
319
352
393
F-Rei1
Flag-
A
B
C
F-Reh1
Flag-
a
b
c
1
358
391
432
F-RR1
Flag-
A
b
c
F-RR2
Flag-
A
B
c
F-RR3
Flag-
a
B
C
F-RR4
Flag-
a
b
C
B
*reh1Δ* strain
Tagged rRNA Level (total RNA)
1.5
1.0
0.5
0.0
A2451U
WT strain
vector
Flag-Reh1
Flag-Reh1ΔC
F-RR1
F-RR2
F-RR3
F-RR4
C
10x
10x
+ vector
+ Reh1
+ Rei1
+ Rei1ΔC
+ RR1
+ RR2
+ RR3
+ RR4
*rei1Δ* strain
23° C
30° C
D
10x
10x
WT
*reh1Δ*
*rei1Δ*
*yvh1Δ*
23° C
3 days
30° C
2 days
WT
*reh1Δ*
*rei1Δ*
*yvh1Δ*
free
40S
60S
80S

Figure S3

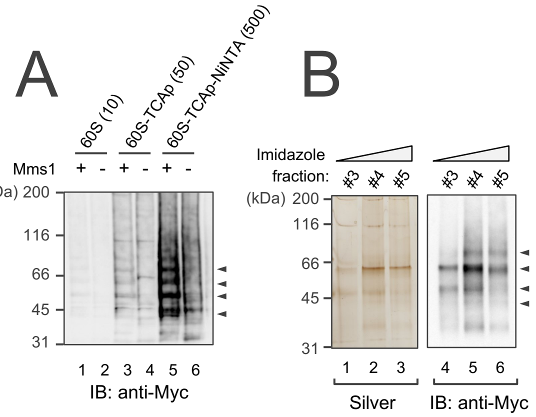


C

MS analysis results:

| | |
|---|---|
| non RPs | Sro9, Hxt5, Ubiquitin |
| RPS | RPS4AB, S7AB, S8AB |
| RPL | RPL12AB, L19AB, L13AB, L20AB, L28 |

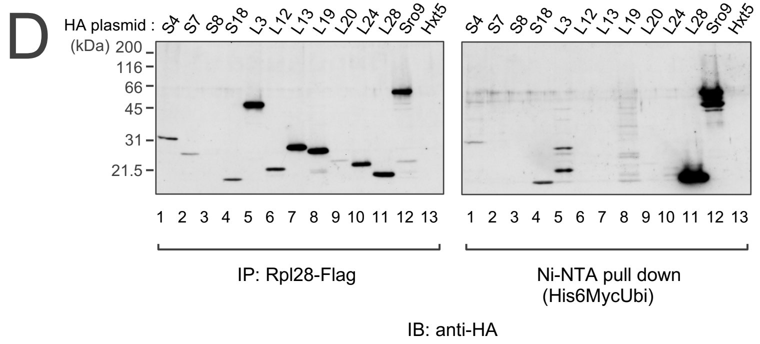


Figure S4

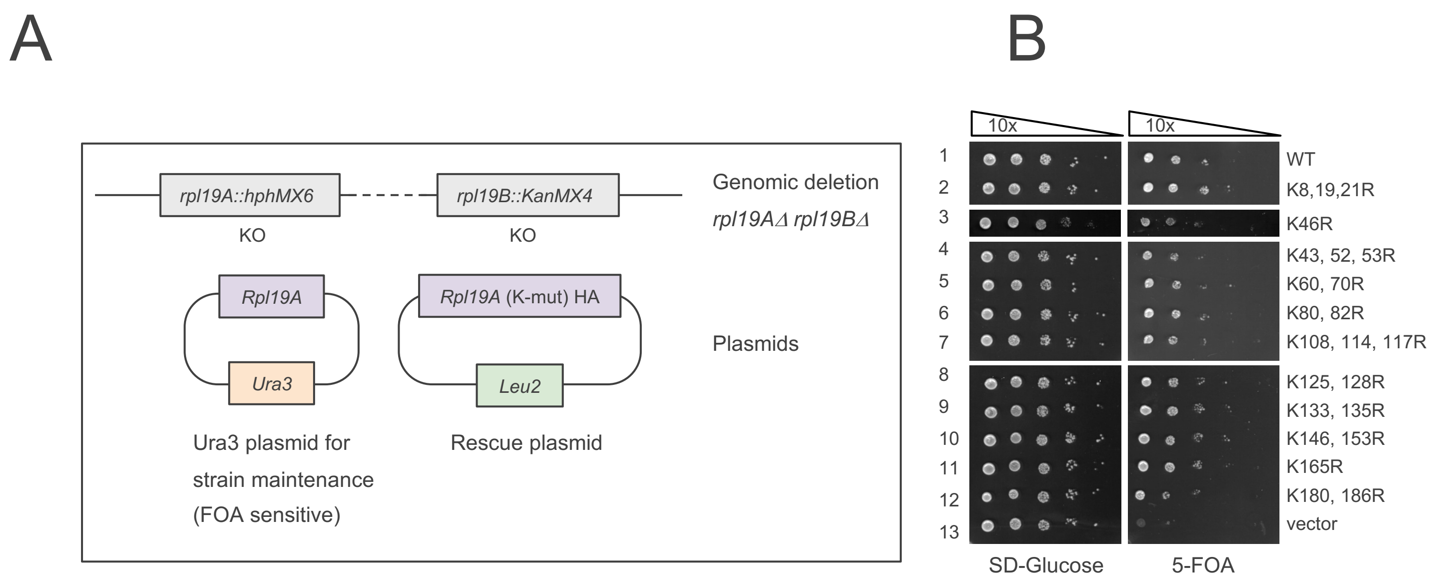
A
B
rpl19A::hphMX6
rpl19B::KanMX4
KO
KO
Genomic deletion
rpl19AΔ rpl19BΔ
Rpl19A
Rpl19A (K-mut) HA
Ura3
Leu2
Plasmids
Ura3 plasmid for
strain maintenance
(FOA sensitive)
Rescue plasmid
10x
10x
1
2
3
4
5
6
7
8
9
10
11
12
13
WT
K8,19,21R
K46R
K43, 52, 53R
K60, 70R
K80, 82R
K108, 114, 117R
K125, 128R
K133, 135R
K146, 153R
K165R
K180, 186R
vector
SD-Glucose
5-FOA

Figure S5

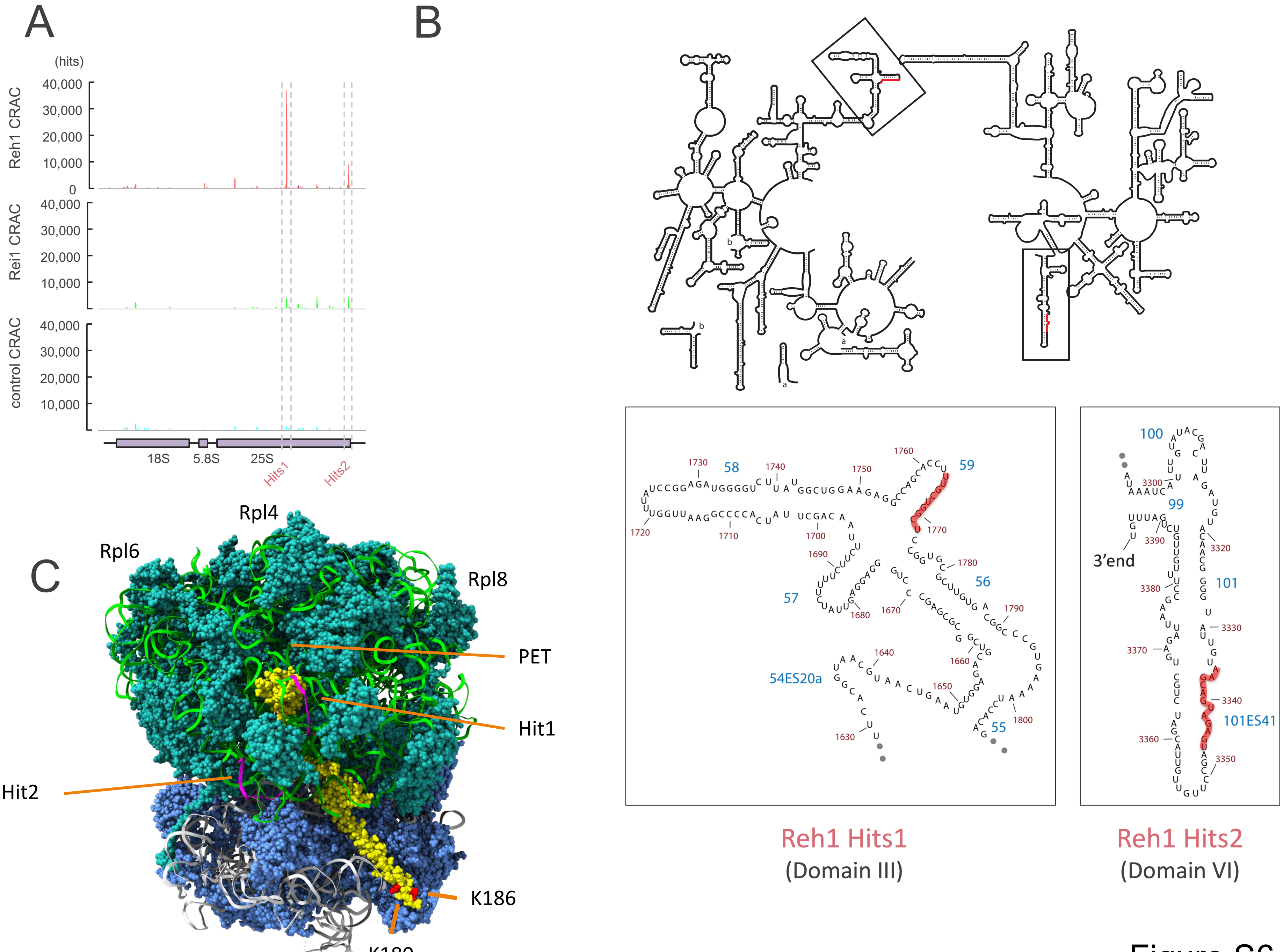

A
B
C
(hits)
Reh1 CRAC
Rei1 CRAC
control CRAC
40,000
30,000
20,000
10,000
0
18S
5.8S
25S
Hits1
Hits2
Rpl4
Rpl6
Rpl8
PET
Hit1
Hit2
K186
K180
58
59
57
56
54ES20a
55
100
99
101
101ES41
3'end
Reh1 Hits1
(Domain III)
Reh1 Hits2
(Domain VI)


Figure S6

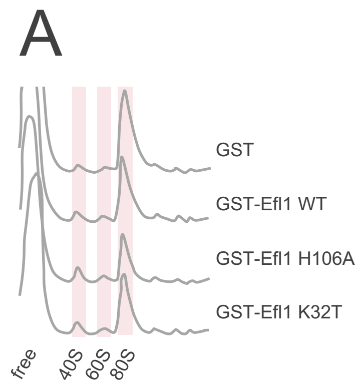
A
GST
GST-Efl1 WT
GST-Efl1 H106A
GST-Efl1 K32T
free
40S
60S
80S

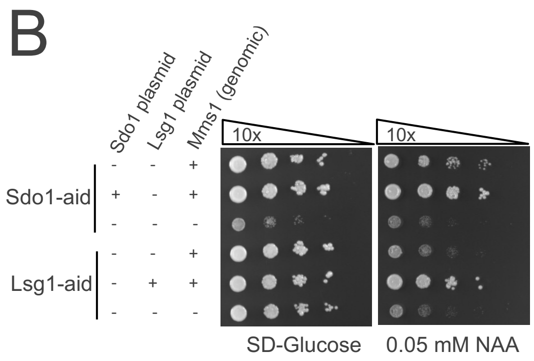
B
Sdo1 plasmid
Lsg1 plasmid
Mms1 (genomic)
Sdo1-aid
Lsg1-aid
10x
10x
SD-Glucose
0.05 mM NAA

Figure S7

### **Table S1** *S. cerevisiae* strains used in this study

| Name | Relevant genotype | Source |
|---|---|---|
| BY4741 | MAT a *hisΔ1 leu2Δ0 met15Δ0 ura3Δ0* | NBRP Yeast, JAPAN |
| BY25598 | MAT a *ura3-1*::ADH1-OsTIR1-9Myc(URA3) *ade2-1 his3-11,15 leu2-3,112 trp1-1 can1-100* | Nishimura *et al.,* 2009 |
| NOY401 | *MAT* a *rpa190-3 ura3 leu2 trp1 can1* | Nogi *et al.,* 1991 |
| MKY17 | *Rpl28*(*Cyh2*)-*Flag* | Fujii *et al.,* 2009 |
| MKY18 | *Rpl28*(*Cyh2*)-*Flag mms1*::kanMX4 | Fujii *et al.,* 2009 |
| MKY23 | *MAT* a *rpa190-3 ura3 leu2 trp1 can1 mms1*::kanMX4 | This study |
| MKY158 | *rpl19a*::hphMX6 *rpl19b*::kanMX4 pRS316-*Rpl19a* | This study |
| MKY425 | *yvh1*::hphMX6 | This study |
| MKY428 | *yvh1*::hphMX6 *crt10*::kanMX4 | This study |
| MKY441 | *yvh1*::hphMX6 *mrt4*::kanMX4 | This study |
| MKY458 | BY25598 *mms1*::hphMX6 | This study |
| MKY461 | BY25598 *Sdo1-aid*::kanMX4 | This study |
| MKY462 | BY25598 *Lsg1-aid*::kanMX4 | This study |
| MKY489 | MKY461 *mms1*::hphMX6 | This study |
| MKY490 | MKY462 *mms1*::hphMX6 | This study |
| MKY510 | *yvh1*::hphMX6 *reh1*::kanMX4 | This study |
| MKY590 | *mex67*::hphMX6 pRS316-*Mex67* | This study |
| MKY591 | *mex67*::hphMX6 *mms1*::kanMX4 | This study |
| MKY592 | *mex67*::hphMX6 *reh1*::kanMX4 | This study |

Other strains, including *mms1Δ*, *paf1Δ*, *hel2Δ*, *rqc1Δ*, *rqc2Δ*, r*ub1Δ*, *lag2Δ*, *yuh1Δ*, *zuo1Δ*, *ssz1Δ*, *arx1Δ*, *reh1Δ*, *rei1Δ* (*slm6Δ*), were obtained from Yeast knock-out MAT a collection (Dharmacon, MAT a kanMX4 *ura3Δ0 leu2Δ0 his3Δ0 met15Δ0*). Each strain was confirmed by genomic PCR analyses before use.

### **Table S2** *S. cerevisiae* plasmids used in this study

| Name | Relevant information | Source |
|---|---|---|
| BYP6998 | pTS910CU-*Reh1-GFP CEN Ura3* | Iwase *et al.,* 2004 |
| pMK004 | gal7 promoter-18S-5.8S-25S (tagged) *2μ Ura3* | Fujii *et al.,* 2009 |
| pMK045 | YCplac111-*Rpl19A*-HA *CEN Leu2* | This study |
| pMK051-063 | pMK045 with mutations (K8, 19, 21R), (K43, 52, 53R), (K60, 70R), (K80, 82R), (K108, 114, 117R), (K125, 128R), (K133, 135R), (K146, 153R), (K165R), (K180, 186R), (K180R), (K186R), (K46R) | This study |
| pMK064 | YCplac111-*Rpl19A CEN Leu2* | This study |
| pMK065-067 | pMK064 with mutation (K180R), (K186R), (K180, 186R) | This study |
| pMK069 | pRS316-*Rpl19A CEN Ura3* | This study |
| pMK088 | pYO323-*Myc-ubiquitin 2μ His3* | Fujii *et al.,* 2009 |
| pMK091 | pYO323-*His6-Myc-Ubiquitin* (G76A) *2μ His3* | This study |
| pMK215 | pRS316-*Mrt4 CEN Ura3* | This study |
| pMK216 | pMK215 with mutation (G68E) | This study |
| pMK222 | YCplac111-*Rpl11A-Flag CEN Leu2* | This study |
| pMK268- 303 | pMK004 with mutation 1-36 (Figure S1) *2μ Ura3* | This study |
| pMK338 | pRS315-Lsg1-HA CEN Leu2 | This study |
| pMK351 | YEplac195-Gal7 promoter-*GST 2μ Ura3* | This study |
| pMK354-356 | pMK351 with *GST-Efl1* wild-type, (H106A), (K32T) | This study |
| pMK361 | pRS315-*Flag-Rei1 CEN Leu2* | This study |
| pMK362 | pRS315-*Flag-Reh1 CEN Leu2* | This study |
| pMK363 | pYO323-Gal1 promoter-*Crt10-HA 2μ His3* | This study |
| pMK364 | BG1805-Gal1 promoter-*Mms1-Flag 2μ Ura3* | This study |
| pMK365 | pYO325-Gal1 promoter-*Rtt101 2μ Leu2* | This study |
| pMK386 | pRS315-*Yvh1-Flag CEN Leu2* | This study |
| pMK380 | YCplac111-*Sdo1-HA CEN Leu2* | This study |
| pMK387,389 | pMK386 with mutation (C117S), ΔN [2-203aa] | This study |
| pMK415 | YCplac111-*Flag-Rei1ΔC*[D352Ter] *CEN leu2* | This study |
| pMK416 | YCplac111-*Flag-Reh1ΔC*[E391Ter] *CEN leu2* | This study |
| pMK417-420 | YCplac111-*Flag-Rei1/Reh1* hybrid RR1-4 (Extended Data Fig. 4b) | This study |
| pMK451 | pRS316-*Flag-His6-Rei1 CEN Ura3* | This study |
| pMK452 | pRS316-*Flag-His6-Reh1 CEN Ura3* | This study |
